# Protection against *APOE4*-associated aging phenotypes with the longevity-promoting intervention 17α-estradiol in male mice

**DOI:** 10.1101/2024.03.12.584678

**Authors:** Cassandra J. McGill, Amy Christensen, Wenjie Qian, Max A. Thorwald, Jose Godoy Lugo, Sara Namvari, Olivia S. White, Caleb E. Finch, Bérénice A. Benayoun, Christian J. Pike

## Abstract

The apolipoprotein ε4 allele (*APOE4*) is associated with decreased longevity, increased vulnerability to age-related declines, and disorders across multiple systems. Interventions that promote healthspan and lifespan represent a promising strategy to attenuate the development of *APOE4*-associated aging phenotypes. Here we studied the ability of the longevity-promoting intervention 17α-estradiol (17αE2) to protect against age-related impairments in *APOE4* versus the predominant *APOE3* genotype using early middle-aged mice with knock-in of human *APOE* alleles. Beginning at age 10 months, male *APOE3* or *APOE4* mice were treated for 20 weeks with 17αE2 or vehicle then compared for indices of aging phenotypes body-wide. Across peripheral and neural measures, *APOE4* was associated with poorer outcomes. Notably, 17αE2 treatment improved outcomes in a genotype-dependent manner favoring *APOE4* mice. These data demonstrate a positive *APOE4* bias in 17αE2-mediated healthspan actions, suggesting that longevity-promoting interventions may be useful in mitigating deleterious age-related risks associated with *APOE4* genotype.

## Introduction

Age is a primary risk factor for systemic impairments, such as metabolic dysfunction and inflammation, as well as neural declines contributing to diminished cognitive abilities and increased risks of Alzheimer’s disease (AD) and related disorders [1]. In human populations, vulnerability to mortality and age-related dysfunction is modulated by genotype at the apolipoprotein E (*APOE*) locus, with generally detrimental effects associated to the ε4 genotype (*APOE4*). Compared to *APOE3*, the most common of *APOE* alleles, *APOE2* is associated with increased longevity and decreased risk of cognitive impairment and AD [2–4]. Conversely, *APOE4* is associated with reduced longevity [2–4] and is the most significant genetic risk factor for both age-related cognitive impairment and AD pathogenesis [3, 5, 6]. The primary roles of apoE protein involve lipid trafficking and cholesterol homeostasis [7–9] through which it impacts metabolism, inflammation, and several other systems [3, 5, 10, 11]. The apoE cascade hypothesis posits that *APOE* genotypes yield variations in apoE structure, lipidation, protein levels, receptor binding, and oligomerization that, in turn, result in diverse functional outcomes [12]. In the context of *APOE4*, these variations are thought to drive body-wide homeostatic alterations that ultimately contribute to numerous age-related declines including vulnerability to cognitive decline and AD [12]. Thus, *APOE4* genotype may be linked to increased risks of dysfunction and disease by regulation of aging processes.

Strategies to improve healthspan and lifespan have notable promise in reducing risks of age-related impairments across multiple body systems, perhaps especially in the context of *APOE4*. A variety of candidate longevity interventions have been tested through the NIA Interventions Testing Program (ITP). Given the relationship between *APOE4* and aging, we hypothesized that longevity interventions may prove efficacious against age-associated *APOE4* phenotypes. Out of the scores of compounds tested by the ITP, 17α-estradiol (17αE2) is one of the few that showed significant improvements in mean and maximum measures of male lifespan [13], as well as improvements in various indices of systemic and brain aging [14–22]. 17αE2 is a naturally occurring diastereomer of the primary estrogen 17β-estradiol but has comparatively weak estrogenic effects owing to reduced affinity for estrogen receptors [23]. 17αE2 has been shown to be protective in brain [24–26] and peripheral tissues [15, 21, 22, 27], in part by improving metabolism and reducing inflammation. The potential efficacy of 17αE2 and other longevity-promoting interventions to attenuate *APOE4* phenotypes associated with deleterious aging outcomes is a promising strategy that has yet to be investigated.

Here, we evaluate the therapeutic potential of 17αE2 to protect against the development of systemic and neural age-related phenotypes in *APOE3* versus *APOE4* genotypes. Specifically, we treated male mice with replacement of mouse *Apoe* with human *APOE3* or *APOE4* during the transition to middle age with 0 or 14ppm 17αE2 for a 20-week period. As expected, the data show age-related phenotypes are exaggerated with *APOE4* genotype. Notably, our findings also demonstrate that 17αE2 treatment leads to improvements in several systemic and brain outcomes more strongly in *APOE4* mice, suggesting a novel strategy to mitigate the progeroid effects of *APOE4* genotype and providing a proof-of-principle of a personalized medicine approach in a preclinical model.

## Results

### 17αE2 improves senescent phenotypes in male APOE4 mice

To determine if the longevity-promoting intervention 17αE2 provides greater protection against systemic aging phenotypes in the context of the *APOE4* allele, we used strains of mice with humanized *APOE3* or *APOE4* sequences [28]. Since the NIA intervention testing program showed anti-aging effects of 17αE2 only in males [13], we decided to focus only on male mice in the study. *Thus*, we fed *APOE3* or *APOE4* knock-in male mice with normal chow or chow supplemented with 14.4ppm 17αE2, the dose shown to have anti-aging effects [29], for a period of 20 weeks starting at 10 months of age (Figure 1A). Even in the absence of AD pathology, *APOE4* is associated with both premature aging phenotypes and decreased lifespan in humans and mice [4]. Thus, we first assessed the ability of 17αE2 to mitigate aging phenotypes using (i) a 25-point frailty index [30] and (ii) a liver DNA methylation-based epigenetic clock [31]. The frailty index measures visible markers of aging including the physical/musculoskeletal system, the ocular/nasal system, the respiratory system, the digestive/urogenital system, and observable signs of discomfort [30]. Notably, we found that control *APOE4* mice have a significantly higher frailty index compared to *APOE3* control, and that *APOE4* mice treated with 17αE2 have a significantly reduced index that is no longer statistically different from control *APOE3* mice (Figure 1B; 2-way ANOVA; F_treatment(1,56)_ = 5.1, *p* = 0.03 ; F_genotype(1,56)_ = 5.1, *p* = 0.03). Consistent with this, using a validated epigenetic clock for mouse aging, we found that *APOE4* animals have a liver DNA methylation signature consistent with that of chronologically relatively older mice, while *APOE3* control and treated mice have a comparatively younger DNA methylation pattern (Figure 1C; 2-way ANOVA; F_genotype(1,36)_ = 12.1, *p* = 0.001). Similar to the frailty index, 17αE2-treated *APOE4* mice were no longer statistically different from *APOE3* controls, exhibiting a DNA methylation pattern “younger” than their chronological age. Together, these data are consistent with the hypothesis that 17αE2 treatment initiated at early middle-age leads to improved aging phenotypes specifically in male *APOE4* mice.

**Figure 1.**
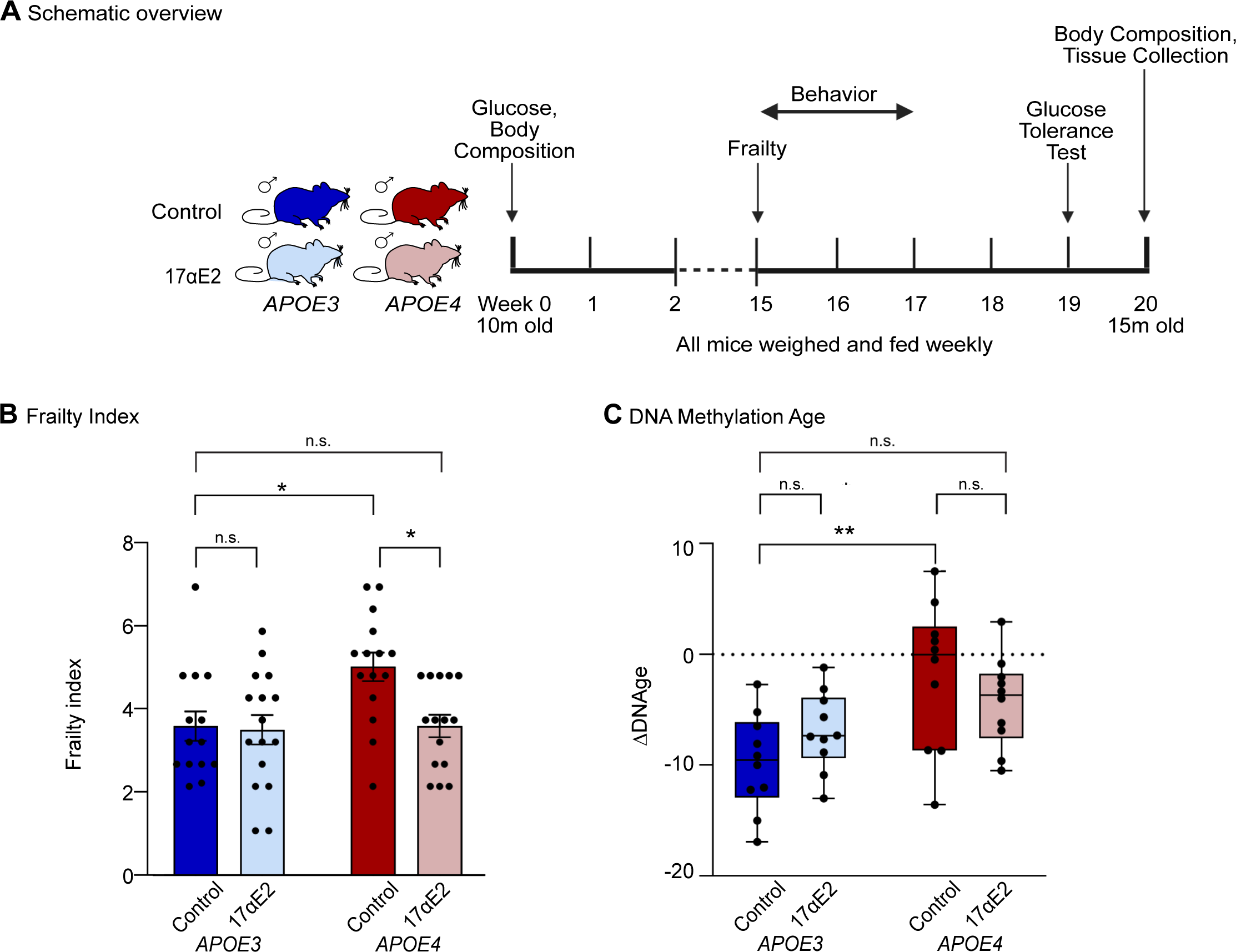
*APOE4* genotype increases aging phenotypes that are ameliorated by 17αE2. (A) Schematic of experimental design. 10 to 10.5-month-old male *APOE3* and *APOE4* targeted replacement mice were fed with normal chow or chow supplemented with 14.4ppm 17αE2 for 20 weeks. At 15 weeks of treatment, animals were assessed for frailty and behavioral changes. At 19 weeks of treatment a glucose tolerance test was performed, and tissues were collected at 20 weeks post treatment initiation (n=19-22/group). (B) Measure of systemic aging using a phenotypic 25-point frailty index (n=14-16/group). (C) Measure of biological aging using a DNA methylation clock derived from liver tissue (n=10/group). In (B) and (C) dark blue indicates *APOE3* control, light blue indicates *APOE3* 17αE2, dark red indicates *APOE4* control, and light red indicates *APOE4* 17αE2. Data show mean ± SEM. Asterisks denote statistical significance: * p < 0.05, *** p < 0.001 in 2-way ANOVA Tukey post-hoc test.

### 17αE2 decreases body weight and food intake more strongly in male APOE4 mice

17αE2 was associated with significant reductions in food intake in both the *APOE3* and *APOE4* mice (Figure 2B, 2-way repeated measures ANOVA; F_treatment(3,83)_ = 12.6, *p*<0.0001) with a statistically nonsignificant trend for a stronger effect in *APOE4* (mean -10.8 ± 4.5% food intake) than *APOE3* (mean -4.1 ± 3.9% food intake) mice. 17αE2 treatment also resulted in significant decreases in body weight (Figure 2C, 2-way repeated measures ANOVA; F_treatment(3,78)_ = 23.1, *p*<0.0001); however, *APOE4* mice on 17αE2 displayed greater reductions (mean -17.2 ± 1.8% body weight,) compared to the *APOE3* 17aE2 treated mice (mean -6.2 ± 0.8% body weight). Note that at week 0 of treatment, *APOE4* mice exhibited significantly higher body weight compared to *APOE3* mice (Supplemental Figure 1A), as well as significantly higher fat mass (Fig. 2D) and lower lean mass (Fig. 2E). Body composition analysis at week 20 revealed a similar pattern of 17αE2 treatment improvements with both genotypes exhibiting trends for reduced relative fat mass (Figure 2D, 3-way ANOVA; F_treatment(1,112)_ = 11.4, *p* = 0.001) and increased relative lean mass (Figure 2E, 3-way ANOVA; F_treatment(1,112)_ = 15.4, *p* = 0.0002). There was a non-significant trend of 17αE2 having a stronger effect in *APOE4* mice, with 17αE2-treated *APOE4* mice having greater reductions of fat mass and more lean mass (Table 1). Importantly, following 17αE2 treatment, *APOE3* and *APOE4* mice did not significantly differ in measures of body morphometry. These findings were paralleled by tissue weights of visceral and retroperitoneal adipose depots, with reductions reaching statistical significance only in the *APOE4*-treated mice (Supplemental Figure 1B-C, 2-way ANOVA; Visceral: F_genotype(1,58)_ = 49.8, *p* < 0.0001; F_treatment(1,58)_ = 18.7, *p* < 0.0001; Retroperitoneal: F_genotype(1,58)_ = 11.7, *p* = 0.001; F_treatment(1,58)_ = 24.8, *p* < 0.0001). Thus, as reported in genetically heterogeneous mice with wild-type murine *Apoe* [22], 17αE2 was associated with reductions in food intake and body weight in mice with knock-in of human *APOE,* though *APOE4* mice experienced relatively greater improvements.

**Figure 2.**
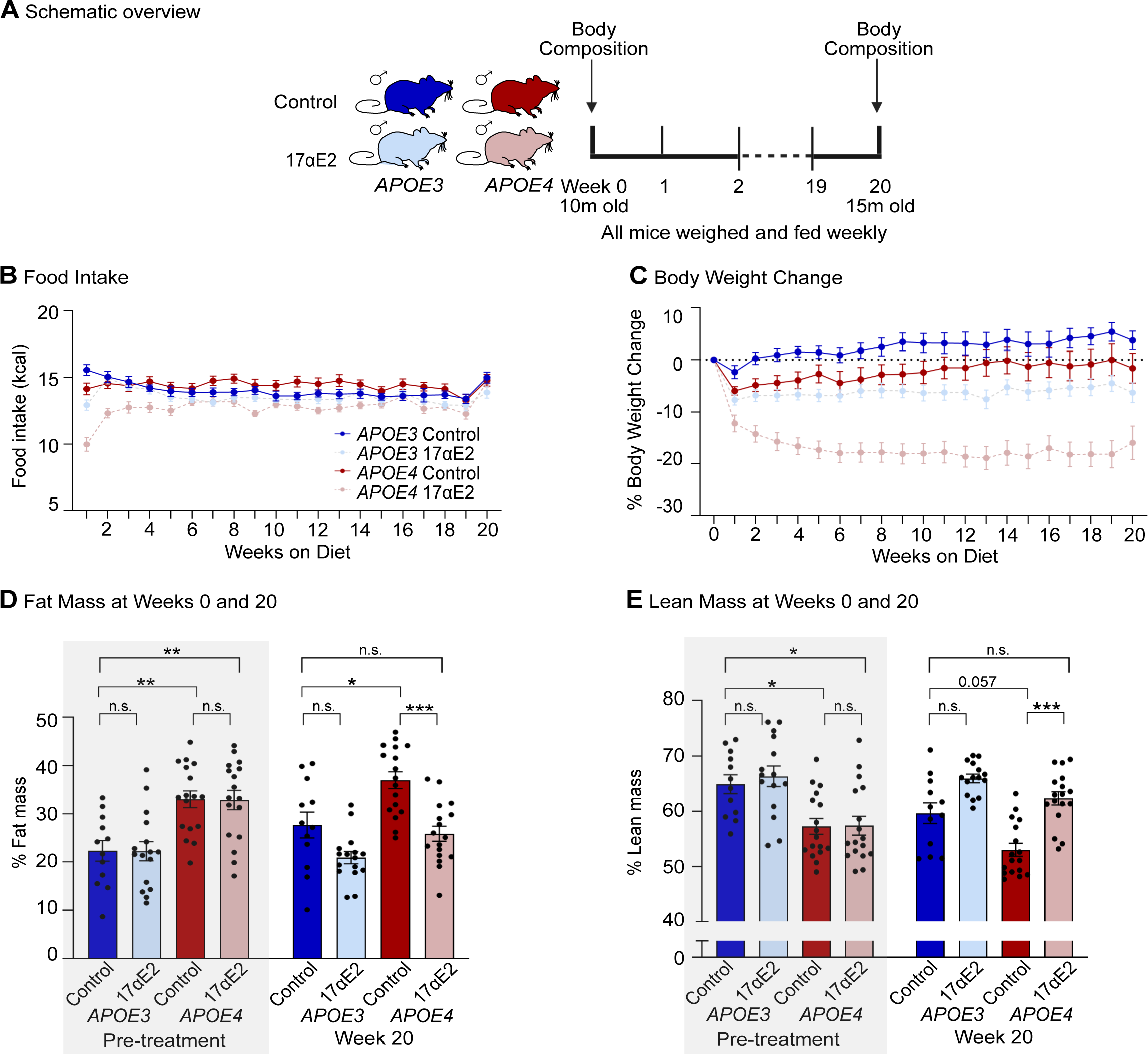
17α estradiol decreases body weight and food intake in middle-aged *APOE3* and *APOE4* targeted replacement mice. (A) Schematic overview. Food intake and body weight were measured weekly. Fat/Lean mass was measured prior to treatment and after 20 weeks of treatment. (B) Food intake (kcal) across the 20 weeks of treatment (n=19-22/group). (C) Percent body weight change across the 20 weeks of treatment (n=19-22/group). (D) Fat mass measured by the Bruker Mini Spec (n=12-17/group). (E) Lean mass measured the Bruker Mini Spec (n=12-17/group). In (D) and (E), shaded panels indicate pre-treatment. In panels (B) and (C), dark blue solid lines indicate *APOE3* control, light blue dashed lines indicate *APOE3* 17αE2, dark red solid lines indicate *APOE4* control, and light red dashed lines indicate *APOE4* 17αE2. In (D) and (E), dark blue indicates *APOE3* control, light blue indicates *APOE3* 17αE2, dark red indicates *APOE4* control, and light red indicates *APOE4* 17αE2. Data show mean ± SEM. Asterisks denote statistical significance: * p < 0.05, ** p < 0.01, *** p < 0.001 in 2-way ANOVA Tukey post-hoc test.

**Table 1.**
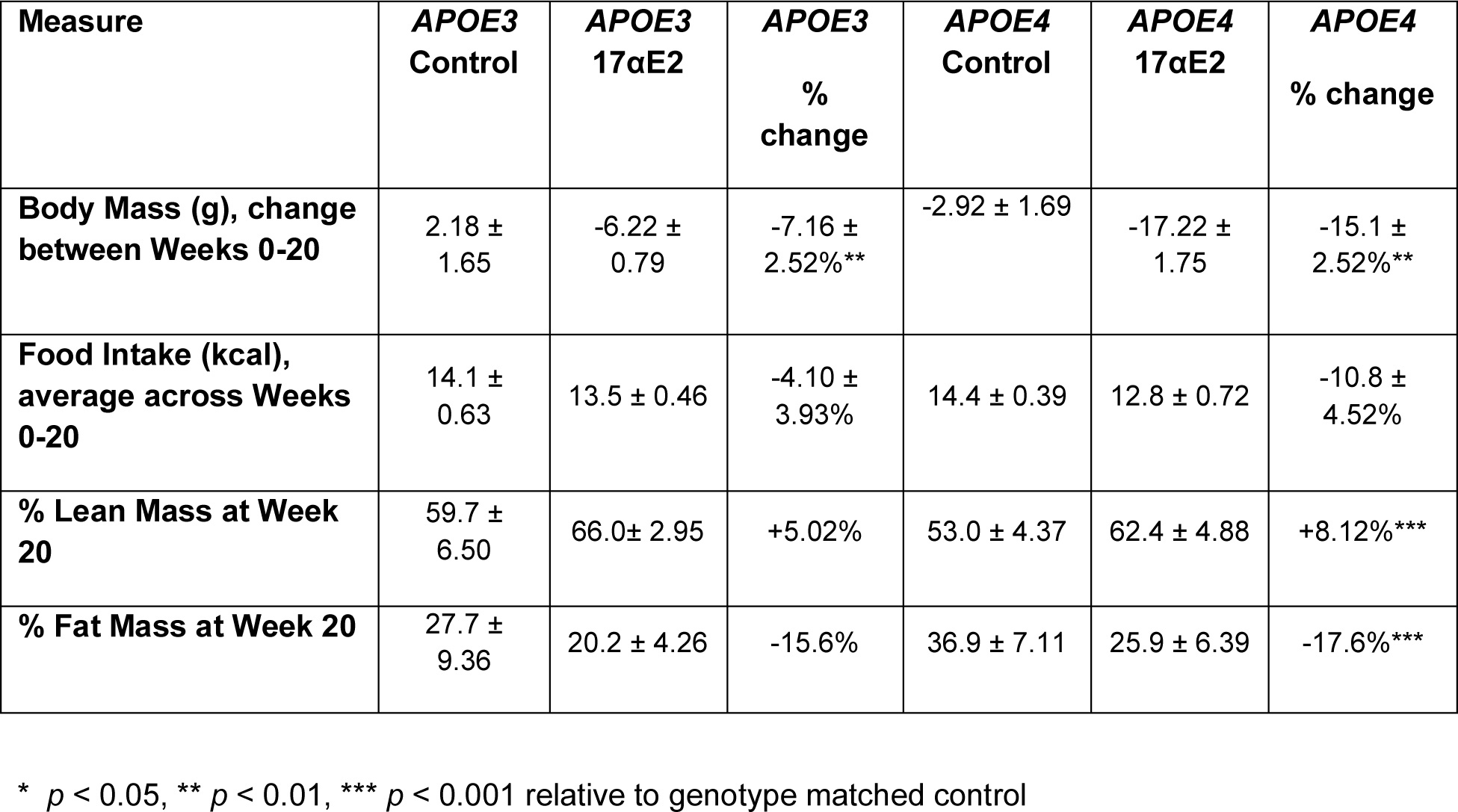
Comparisons of 17αE2-induced changes in body weight, food intake, and lean and fat masses in *APOE3* versus *APOE4* mice.

| Measure | *APOE3*<br>Control | *APOE3*<br>17αE2 | *APOE3*<br>%<br>change | *APOE4*<br>Control | *APOE4*<br>17αE2 | *APOE4*<br>% change |
| --- | --- | --- | --- | --- | --- | --- |
| **Body Mass (g), change between Weeks 0-20** | 2.18 ± 1.65 | -6.22 ± 0.79 | -7.16 ± 2.52%** | -2.92 ± 1.69 | -17.22 ± 1.75 | -15.1 ± 2.52%** |
| **Food Intake (kcal), average across Weeks 0-20** | 14.1 ± 0.63 | 13.5 ± 0.46 | -4.10 ± 3.93% | 14.4 ± 0.39 | 12.8 ± 0.72 | -10.8 ± 4.52% |
| **% Lean Mass at Week 20** | 59.7 ± 6.50 | 66.0 ± 2.95 | +5.02% | 53.0 ± 4.37 | 62.4 ± 4.88 | +8.12%*** |
| **% Fat Mass at Week 20** | 27.7 ± 9.36 | 20.2 ± 4.26 | -15.6% | 36.9 ± 7.11 | 25.9 ± 6.39 | -17.6%*** |

\* $p < 0.05$ , \*\* $p < 0.01$ , \*\*\* $p < 0.001$ relative to genotype matched control

### 17αE2 improves metabolic measures more strongly in APOE4 mice

Consistent with the higher adiposity in *APOE4* mice, we found significantly greater hepatic steatosis in control animals with the *APOE4* vs *APOE3* genotype (Figure 3B, Mann-Whitney test, *p* = 0.02). There is only a significant reduction of hepatic steatosis by 17αE2 treatment in *APOE4* mice (Figure 3B, Mann-Whitney test, *p* = 0.005). In addition to increased fatty liver, *APOE4* control mice exhibited decreased glucose tolerance and increased plasma leptin (Figure 3C-E). In the glucose tolerance test, both *APOE3* and *APOE4* mice treated with 17αE2 displayed significant improvement (Figure 2D, 2-way ANOVA; F_genotype(1,55)_ = 11.6, *p* = 0.001; F_treatment(1,55)_ = 24.2, *p* < 0.0001). Similar to the outcome of the liver lipid staining, plasma leptin levels were significantly decreased only in the *APOE4* treated animals. (Figure 2E, 2-way ANOVA; F_genotype(1,42)_ = 10.2, *p* = 0.003; F_treatment(1,42)_ = 15.4, *p* = 0.0003). Thus, our data show that 17αE2 treatment improved metabolic phenotypes in both male *APOE3* and *APOE4* mice, with stronger benefits generally observed in *APOE4* mice.

**Figure 3.**
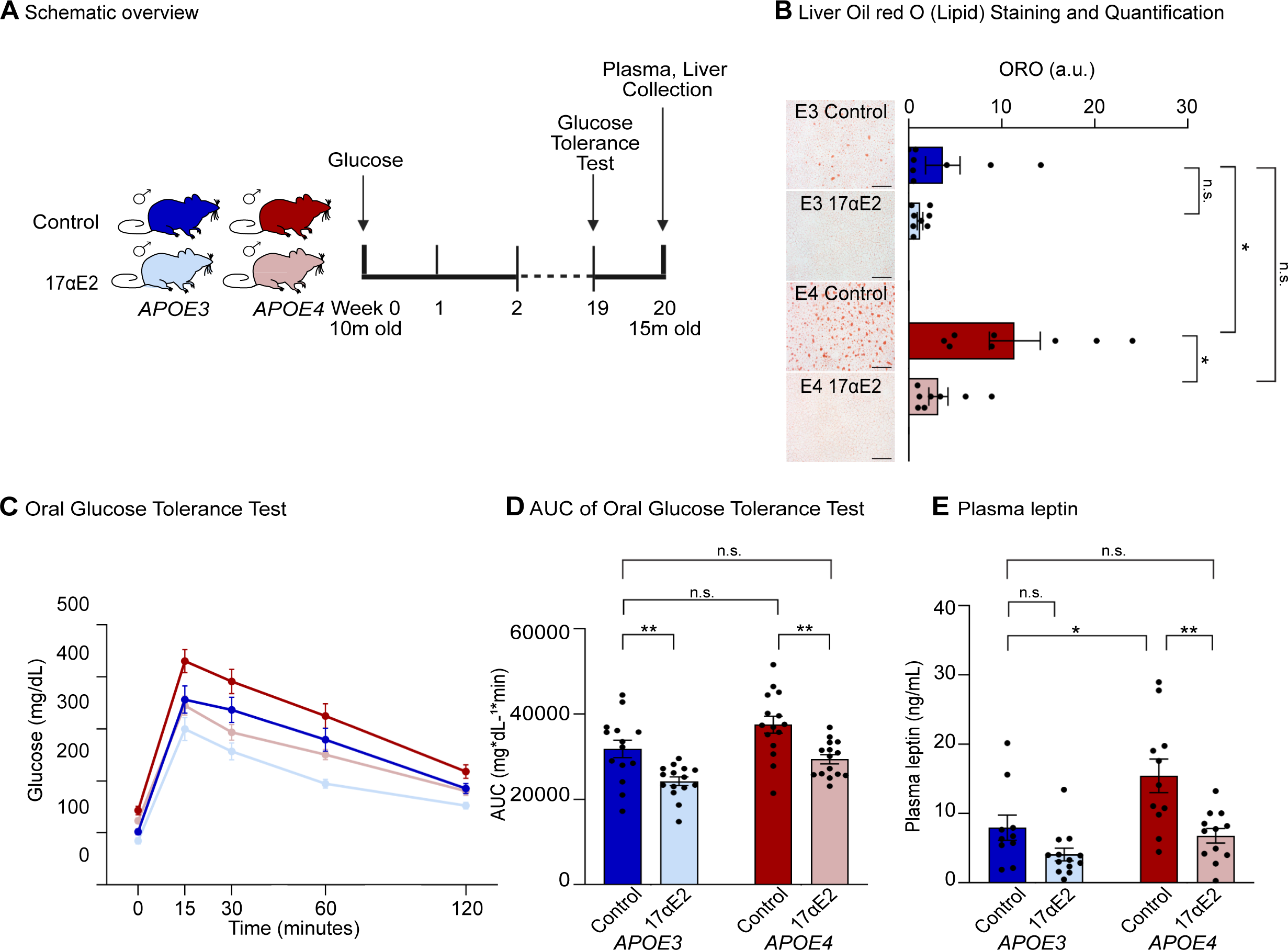
*APOE4* mice have greater metabolic dysfunction and greater protection with 17αE2 compared to *APOE3* mice. (A) Schematic overview. Glucose tolerance test was performed after 19 weeks of treatment and tissues were collected after 20 weeks. (B) ORO quantification (n=8/group). Images show oil red O labeling (ORO) of lipid accumulation in livers of 15-month-old male *APOE3* and *APOE4* mice treated with 0 or 14.4 ppm 17αE2. Scale bar size indicates 100µM. (C) Oral glucose tolerance test (GTT) measured at baseline then 15-, 30-, 60-, and 120-minutes post oral gavage with glucose. (D) Area under the curve analysis for GTT seen in (D) (n=14-15/group). (E) Plasma leptin measured through ELISA (n=12-13/group). Data show mean ± SEM. In (B) through (E), dark blue indicates *APOE3* control, light blue indicates *APOE3* 17αE2, dark red indicates *APOE4* control, and light red indicates *APOE4* 17αE2. Asterisks denote statistical significance: * p < 0.05, ** p < 0.01, *** p < 0.001, **** p < 0.0001 in Mann Whitney test for ORO, 2-way ANOVA Tukey post-hoc test for all other analyses.

### 17αE2 reduces genotype-specific differences between APOE3 and APOE4 mice

As apoE is a key regulator of the lipidome, we next performed shotgun lipidomics on animals from all groups (Figure 4A). Given the body-wide effects of *APOE*, we analyzed both peripheral (plasma) and neural (cerebral cortex) tissues.

**Figure 4.**
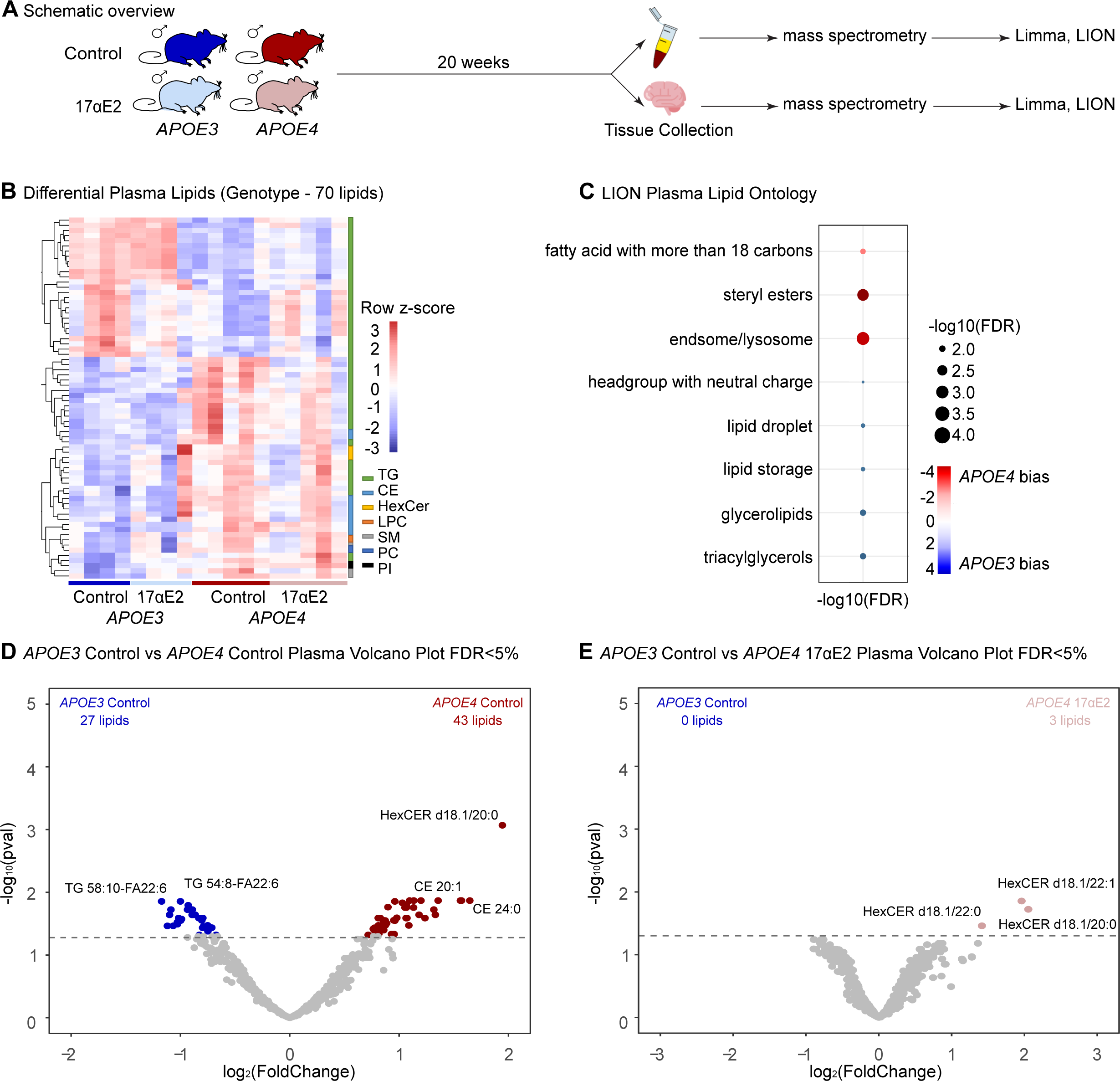
17αE2 reduces genotype-specific differences in the plasma lipidome of *APOE3* and *APOE4* mice. (A) Schematic overview. Plasma and cortex were isolated from the RNA-seq cohort after 20 weeks of treatment. (B) Differentially abundant lipids between *APOE3* and *APOE4* control mice plasma (FDR 5%) (n=4-5/group) Red indicates higher abundance and blue indicates lower abundance. (C) LION lipid ontology analysis top enriched pathways from the 70 lipids seen in (B). Red indicates *APOE4* bias in plasma, blue indicates *APOE3* bias in plasma. (D) Volcano plot showing differentially abundant lipids between *APOE3* and *APOE4* control mice plasma (FDR 5%). (E) Volcano plot showing differentially abundant lipids between *APOE3* control and *APOE4* 17αE2 mice plasma (FDR 5%).

In the plasma (at 15 months old), we identified 70 lipids with significant differential abundances between *APOE3* and *APOE4* control mice (Figure 4B,D, Supplemental Table 1A, FDR < 0.05). This list consists of triglycerides (TG), hexosylceramides (HexCER), and sphingomyelin (SM), among others. Enrichment analysis using LION [32] revealed that lipids upregulated in the plasma of *APOE4* are primarily involved with endosome/lysosomes and steryl esters, while *APOE3* plasma is enriched for lipids involved in lipid storage (Figure 4C, Supplemental Table 1C). To our knowledge, there are currently no other plasma shotgun lipidomics datasets reported from *APOE* knock-in mouse models. Comparing to previous findings in humans, several lipid types associated with both *APOE4* and AD were also significantly differentially regulated in our dataset. Plasma ceramides (CE) CE (17:0), CE (18:0), CE (20:1), CE (22:4), lysophosphatidylcholine (LPC) (20:0), and phosphatidylinositol (PI) (18:0/20:3) were found [33] to be increased in individuals with AD. CE (22:4), CE (24:1), and HexCER (d18:1/22:0) were found to be positively correlated with both *APOE4* and AD [34]. In these prior reports [33, 34], CE (22:4) was found to be positively correlated with AD. Indeed, these AD-associated lipids were all found to be increased in *APOE4* control mice. However, between the control *APOE3* and 17αE2-treated *APOE4* groups there were only three lipids with significant differential abundance, all of which are HexCERs (Figure 4E, Supplemental Figure 2C-E, Supplemental Table 1B, FDR < 0.05). Between *APOE3* control and treated groups, as well as between *APOE4* control and treated, no significantly different lipids were identified, likely because of tissue-specific differences in 17αE2 actions. Finally, an analysis at the level of overall lipid classes revealed a significant interaction between genotype and treatment in CE, with CE being increased with 17αE2 treatment only in *APOE3* mice (Supplemental Figure 2A; 2-way ANOVA; F_interaction(1,14)_ = 7.5, *p* = 0.02; F_treatment(1,14)_ = 8.1, *p* = 0.01). There are significant main effects of both genotype and treatment in HexCER abundance (Supplemental Figure 2A; 2-way ANOVA; F_genotype(1,14)_ = 9.5, *p* = 0.008; F_treatment(1,14)_ = 11.71, *p* = 0.004). In humans, sphingolipid dysregulation occurs with aging and AD, specifically increases in long-chain ceramides including HexCer [35]. Interestingly, increased levels of total HexCer have been reported in a centenarian population [36]. There was also a significant interaction between genotype and treatment in phosphatidic acid (PA), with 17αE2 slightly increasing PA in *APOE3* treated mice, while decreasing PA in *APOE4* 17αE2-treated mice (Supplemental Figure 2A; 2-way ANOVA; F_interaction(1,14)_ = 5.8, *p* = 0.03). Lastly, there was an overall effect of treatment on phosphatidylcholine (PC) abundance, with 17αE2 increasing PC in both *APOE3* and *APOE4* mice (Supplemental Figure 2A; 2-way ANOVA; F_treatment(1,14)_ = 4.8, *p* = 0.05). Taken together, while 17αE2 did not significantly change the individual lipid species content between control and treated groups within *APOE* genotypes, it did significantly diminish the genotype differences found between *APOE3* and *APOE4* mice.

In contrast to observations in plasma, we did not find any significant differences in individual lipids based on genotype or 17αE2 treatment in cerebral cortex tissue. Interestingly, we did find significant *APOE* genotype differences in overall lipid classes abundances. Specifically, HexCERs are significantly increased in *APOE4* controls compared to *APOE3* controls (Supplemental Figure 2B, 2-way ANOVA; F_genotype(1,15)_ = 10.4, *p* = 0.006). Previous studies have assessed the cortical lipidome in *APOE4* mice using a targeted approach [37] and, similar to our dataset, found increased amounts of total HexCer in the cortex of *APOE4* homozygous mice relative to *APOE3* mice [37]. PA levels are decreased in human *APOE4* AD patients, and phospholipid dysregulation is strongly implicated in *APOE4*-associated AD pathogenesis [38–40]. PA was also increased in both *APOE4* groups compared to *APOE3* (Supplemental Figure 2B, 2-way ANOVA; F_genotype(1,15)_ = 9.5, *p* = 0.008). Sphingomyelin dysregulation is associated with *APOE4* and AD [35, 37, 41]. Interestingly, sphingomyelin was found to be increased in both groups treated with 17αE2 (Supplemental Figure 2B, 2-way ANOVA; F_treatment(1,15)_ = 6, *p* = 0.03).

Collectively, our lipidomic data show that 17αE2 treatment in male mice dampens *APOE* genotype-specific differences in the plasma lipidome, with more limited effects in the cerebral cortex.

### 17αE2 reduces genotype-specific differences between APOE3 and APOE4 microglia

*APOE* genotype has established effects on innate immunity and microglia [42–44], both of which are also linked to increased risks of cognitive decline and AD. To understand how microglia respond to systemic 17αE2 treatment in the context of *APOE3* vs. *APOE4* genotypes, we isolated primary microglia from 15-month-old male treated and control mice using magnetic-activated cell sorting (MACS) technology (Figure 5A). Importantly, these mice were not subjected to any behavioral or metabolic assessments prior to euthanasia, and were processed together to minimize undesired batch effects [45]. Transcriptomes of isolated primary microglia were profiled by RNA-sequencing (RNA-seq) (Figure 5A).

**Figure 5.**
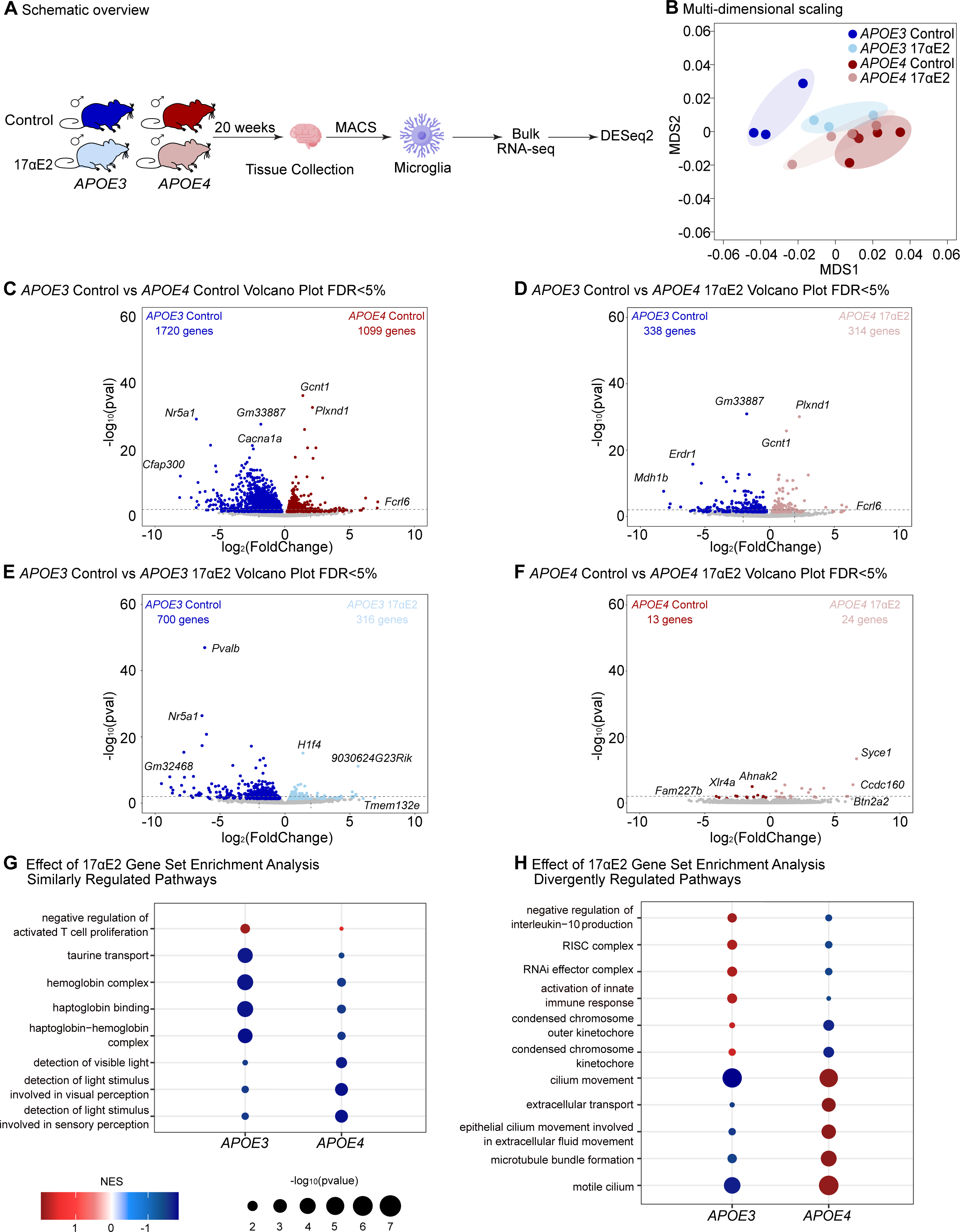
17αE2 reduces genotype-specific differences in the microglia of *APOE3* and *APOE4* mice. (A) Schematic overview. Microglia were isolated from the RNA-seq cohort after 20 weeks of treatment. (B) Multi-dimensional scaling (MDS) of transcriptomes for microglia from all four groups (n=3-5/group). (C) Volcano plot of differentially expressed genes (DESeq2 FDR < 5%) between *APOE3* control (dark blue) and *APOE4* control (dark red) microglia. (D) Volcano plot of differentially expressed genes (FDR < 5%) between *APOE3* control (dark blue) and *APOE4* 17αE2 (light red) microglia. (E) Volcano plot of differentially expressed genes (FDR < 5%) between *APOE3* control (dark blue) and *APOE3* 17αE2 (light blue) microglia. (F) Volcano plot of differentially expressed genes (FDR < 5%) between *APOE4* control (dark red) and *APOE4* 17αE2 (light red) microglia. (G) Effect of 17αE2 in *APOE3* and *APOE4* microglia gene set enrichment analysis (gene ontology). Bubble plot shows top similarly changed gene sets (DESeq2 FDR < 5%). (H) Effect of 17αE2 in *APOE3* and *APOE4* microglia gene set enrichment analysis (gene ontology). Bubble plot shows top divergently changed gene sets (DESeq2 FDR < 5%).

To first assess the similarity of the microglial transcriptomes from our different groups of mice, we utilized multidimensional scaling (MDS) [46]. Interestingly, MDS analysis showed clear separation of samples by genotype and treatment (Figure 5B). Notably, the greatest separation was seen between *APOE3* and *APOE4* control samples, while the 17αE2-treated groups tended to cluster closer together (Figure 5B). To understand the impact of *APOE* genotype and 17αE2 treatment on middle-aged male microglia, we performed differential gene expression analysis using DESeq2 to reveal transcriptional features with significant genotype- or treatment-related regulation at False Discovery Rate (FDR) < 5% using multivariate linear modeling (Figure 5C-F, Supplementary table 2A-D). Importantly, we checked the quality of our dataset for appropriate expression of microglia-specific genes, and lack of expression of other common brain cell types (Supplemental Figure 3A). There were 2,819 genes significantly differentially expressed between control *APOE3* and *APOE4* microglia; however, when comparing control *APOE3* with 17αE2 *APOE4* microglia, only 652 genes were differentially expressed. When comparing *APOE3* control and treated groups, there were 1,016 differentially expressed genes. Only 37 differentially expressed genes were identified between control and treated *APOE4* groups, suggesting a larger effect on the microglia transcriptome in the *APOE3* animals. Interestingly, comparison of transcriptome-wide changes showed that 17αE2 treatment in *APOE3* mice is associated with acquisition of a more *APOE4*-like transcriptional landscape, with very few genes changing in the same direction across both genotypes (Supplemental Figure 3B,C).

Next, we asked which functional Gene Ontology gene sets were regulated by 17αE2 treatment within the *APOE3* and *APOE4* groups. Looking at the top 5 up- and downregulated significantly changed gene sets per genotype upon 17αE2 treatment, we found gene sets both similarly and divergently regulated by 17αE2 in the *APOE3* and *APOE4* groups. Sets similarly regulated by 17αE2 included those relating to sensory reception, “haptoglobin-hemoglobin complex”, “negative regulation of activated T cell proliferation”, and “taurine transport”. Haptoglobin and hemoglobin have both been implicated in microglia-related inflammation, with studies suggesting haptoglobin regulates microglia-induced inflammation [47] Intriguingly, a recent study reported that taurine is crucial to healthy aging, and deficiency drives aging phenotypes [48] (Figure 5G). Divergently regulated gene sets included “cilium movement”, “microtubule bundle formation”, and “extracellular transport”, with these being upregulated in the *APOE4* 17αE2-treated group (Figure 5H). The *APOE3* 17αE2-treated group showed upregulation of gene sets relating to “condensed chromosome outer kinetochore”, “activation of innate immune-response”, “RISC complex”, “RNAi effector complex”, and “negative regulation of interleukin-10 production” (Figure 5H). Thus, while 17αE2 treatment appears to be affecting the microglia transcriptome differently between *APOE3* and *APOE4* mice, both genotypes exhibit changes in gene sets relating to metabolic/immune system processes.

### 17αE2 improves brain phenotypes specifically in APOE4 mice

After 15 weeks of 17αE2 treatment, animals from all groups were subjected to behavioral assessments. To measure effects of *APOE* and 17αE2 treatment on general measures of motor activity and anxiety, mice were assessed in the open field task. There were no significant differences across the groups in the time spent exploring the center of the field (Supplemental Figure 4A) although there was a significant genotype effect in the total distance traveled with *APOE4* animals showing higher levels (Supplemental Figure 4B). There were no significant main effects of genotype or treatment on behavioral performance in tasks of exploratory behavior and short-term spatial memory (spontaneous alternation behavior in the Y-maze; Supplemental Figure 4C) and episodic and recognition memory (novel object placement and recognition; Supplemental Figure 4D, E).

In the Barnes maze test of spatial learning and memory, we observed *APOE4*-associated deficits, consistent with previous findings from other groups [49]. In this task, evidence of learning over successive days of trials is indicated by shorter latencies and path lengths in reaching the target as well as reductions in the number of errors in choosing the target. Escape latencies showed significant main effects for time (Figure 6B, 3-way ANOVA; F_time(2.5, 116.1)_ = 8.02, *p* = 0.0002) and genotype (Figure 6B, 3-way ANOVA; F_time(1,46)_ = 4.06, *p* = 0.05), which indicate learning by all groups but poorer performance by *APOE4* mice (Figure 6B). There were significant reductions in path lengths across groups (Figure 6C, 3-way ANOVA; F_time(2.6, 119)_ = 10.4, *p* < 0.0001), consistent with learning. *APOE4* mice showed a nonsignificant trend of shorter path lengths, however relative levels of reduction over the training days (i.e., learning) trended in favor of *APOE3* mice (control: 20.4% ± 9.1, 17αE2: 17.4% ± 7.7 reductions) versus *APOE4* mice (control: 10.3% ± 6.5, 17αE2: 5.2%± 3.7 reductions). Neither latency nor path length were significantly impacted by 17αE2 treatment. Error number showed a significant interaction between time and *APOE* genotype (Figure 6D, 3-way ANOVA; F_timexgenotype(3,138)_ = 7.8, *p* < 0.0001). Specifically, both *APOE3* groups had decreased errors over days, whereas *APOE4* mice showed divergent patterns with performance slightly worsening in *APOE4* control mice and improving in *APOE4* treated mice (Figure 6D). Error number showed a similar pattern on the probe trial: significantly more errors were observed in control but not treated *APOE4* mice relative to control *APOE3* mice (Figure 6E, 2-way ANOVA Tukey’s multiple comparisons; *p* = 0.01). Together, the Barnes maze data are consistent with prior reports of deficits in spatial learning and memory performance in *APOE4* mice [10, 50] and suggest partial functional improvements with 17αE2 treatment in terms of error level.

**Figure 6.**
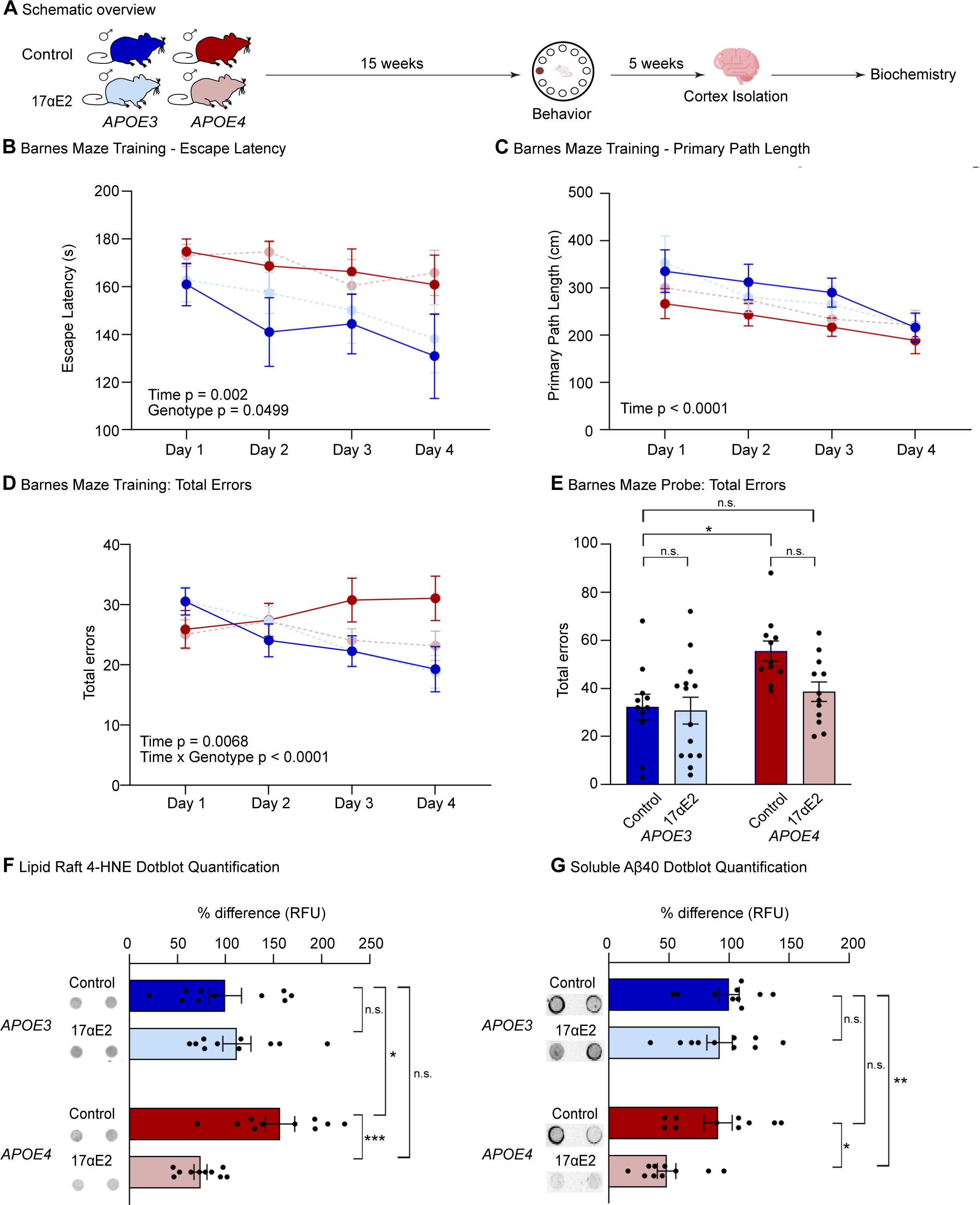
*APOE4* mice have greater neural benefits from 17αE2 than *APOE3* mice. (A) Schematic overview. Animals were subjected to behavior tests after 15 weeks of treatment. One hemibrain was fixed for immunohistochemistry and the other the cortex was isolated for protein extraction. (B) Escape latency during the Barnes maze training days (n=11-14/group). (C) Primary path length to exit hole during the Barnes maze training days (n=11-14/group). (D) Total errors during the Barnes maze training days (n=11-14/group). (E) Total errors during the Barnes maze probe trial (n=11-14/group). (F) Lipid Raft 4-HNE dot blot quantification (n=10/group). (G) Soluble Aβ40 quantification in homogenates of cerebral cortex (n=10/group). In (B) through (G), dark blue indicates *APOE3* control, light blue indicates *APOE3* 17αE2, dark red indicates *APOE4* control, and light red indicates *APOE4* 17αE2. Data show mean ± SEM. Asterisks denote statistical significance: * p < 0.05, ** p < 0.01, *** p < 0.001 in 2-way ANOVA Tukey post-hoc test.

As both *APOE* [12, 38, 50–54] and 17αE2 [14, 17–19, 24] exert a range of effects on the brain, we also considered additional neural outcomes. First, we considered measures of gliosis. We quantified immunohistochemical burden of markers for both astrogliosis (glial fibrillary acid protein) and microgliosis (ionized calcium binding adaptor molecule 1), established measures of age-related glial activation [55, 56]. In the CA1 region of the hippocampus, we observed a modest but significant effect of genotype on astrogliosis, with *APOE4* mice having higher levels than *APOE3* mice (Supplemental Figure 4F). There is a non-significant trend for an overall effect of treatment to reduce astroglia load (p = 0.09) (Supplemental Figure 4F). We found no significant differences in microglial burden across the groups (Supplemental Figure 4G).

Next, we examined levels of lipid peroxidation through a product of oxidized lipids, 4 - hydroxy-2-nonenal (HNE). HNE is a protein carbonyl that forms adducts of cysteine, lysine, or histidine residues and is considered a stable marker of oxidative damage [57]. Lipid peroxidation is observed with normal brain aging and further in AD [57–60] and is particularly strong in lipid rafts. Indeed, recent evidence in human samples shows that 4HNE levels in cerebrocortical lipid rafts are robustly elevated by both *APOE4* and AD [60]. Consistent with these data, we see significantly higher 4-HNE in cerebrocortical lipid rafts from *APOE4* control mice relative to the *APOE3* controls (Figure 6F, 2-way ANOVA Tukey’s multiple comparisons; *p* = 0.03). Notably, there was a significant treatment X genotype interaction (Figure 6F, 2-way ANOVA; F_interaction(1,36)_ = 11.6, *p* = 0.002) in which 17αE2 treatment induced significant decreases in 4-HNE levels in *APOE4* but not *APOE3* mice. Given that lipid rafts are also a key site for generation of amyloid-beta (Aβ) [61], a protein central to AD pathogenesis, we also measured levels of Aβ40 peptides in whole brain homogenates by dot blot technique [61]. We found that 17αE2 treatment was associated with a significant reduction in cerebrocortical soluble Aβ40 in *APOE4* but not *APOE3* mice (Figure 6G, 2-way ANOVA; F_treatment(1,36)_ = 6.6, *p* = 0.01, F_genotype(1,36)_ = 7.4, *p* = 0.01). Taken together, we see that *APOE4* genotype is associated with select deficits in cognition, increased gliosis, and increased lipid peroxidation, which are partially attenuated by 17αE2 treatment.

## Discussion

*APOE* genotype is associated with body-wide changes in lipids, inflammatory tone, and metabolism that are widely posited to underlie its relationships with increased risks of mortality, cognitive impairment, and AD. Here we investigate the hypothesis that interventions established to promote healthspan and longevity may be particularly efficacious against the systemic progeroid phenotypes of *APOE4*. Our findings demonstrate that treatment with the longevity-promoting compound 17αE2 during early middle-age protects against the development of aging phenotypes in an *APOE* genotype-specific manner. Across aging, metabolic, and behavioral measures, we find a similar trend: *APOE4* mice are impaired compared to *APOE3*, and this difference is largely attenuated by 17αE2 such that treated *APOE4* mice appear similar to untreated *APOE3* mice. This was consistent when looking at morphological and epigenetic markers of aging (Figure 1), body mass composition (Figure 2), hepatic steatosis, plasma leptin (Figure 3), the Barnes maze behavioral task, and brain lipid peroxidation (Figure 6). 17αE2-induced improvements in metabolic measures in aging male mice were previously reported [21, 22, 27], but our findings are the first to demonstrate that these improvements are significantly influenced by *APOE* genotype, a genetic factor associated with aging, longevity, and AD [2, 4, 12]. 17αE2-treated *APOE4* mice had improvements in all metabolic parameters measured. In the Barnes maze task of spatial learning and memory, 17αE2-treated *APOE4* mice significantly reduced their total errors, making them statistically indistinguishable from the *APOE3* mice. 17αE2 treatment in *APOE4* mice was also associated with significantly lower lipid raft levels of Aß, a protein implicated as a central factor in AD pathogenesis. Collectively, these findings suggest 17αE2 treatment could be a potential therapeutic against *APOE4*-associated aging phenotypes including increased vulnerability to age-related cognitive decline and AD.

Given the robust evidence that longevity interventions can slow declines in tissue function associated with normal aging that begins in early adulthood and becomes more pronounced by midlife [62, 63], they hold significant promise as protective measures against numerous age-related diseases. Notably, many longevity interventions not only extend lifespan, but also lessen metabolic dysfunction, indicating a complex interplay between longevity and metabolic integrity [13, 22, 29]. Metabolic dysfunction and diseases such as diabetes and hypertension are known drivers of aging phenotypes, cognitive decline, and AD, acting through mechanisms including systemic inflammation [64, 65]. *APOE4* is linked to metabolic dysfunction [5, 10, 11, 40], which may contribute to its relationships with aging processes and age-related diseases. Older adults with *APOE4* are more likely to experience brain glucose hypometabolism, atherosclerosis, and ischemic heart disease [66]. *APOE4* genotype significantly impacts longevity, with most [4, 67, 68] but not all [69] studies associating *APOE4* with increased mortality. Our findings, consistent with existing research, support the notion that *APOE4* induces “pro-aging” phenotypes in both humans and mice [4, 67, 68]. Given that *APOE4* carrier status is observed in ∼15% of the U.S. population [50] and in ∼60% of AD patients [6], there is significant need to identify interventions that can mitigate the heightened risks associated with *APOE4.* This study focuses on 17αE2 as a proof-of-principle intervention, leveraging its well-established multi-system protective effects. We sought to explore 17αE2’s potential in addressing the multifaceted challenges posed by *APOE4*, offering insights for intervention that may extend beyond metabolic health to encompass neuroprotection.

Our research, in conjunction with other studies [13, 29], highlights that longevity drugs are not a “one-size-fits-all” approach to treating age-related impairments and reducing risks for disease. Indeed, a variety of factors interplay in drug responsiveness including age, genetics, and sex dimorphisms. Our specific findings on 17αE2 reveal differential impact in *APOE3* and *APOE4* male mice, with significant protection in *APOE4* mice but a slightly beneficial to neutral effect in *APOE3* male mice. This genotype-dependent effect suggests that the progeroid phenotypes associated with *APOE4* may be especially responsive to longevity-related interventions. Alternative strategies may prove more effective in an *APOE3* context, highlighting the importance of exploring diverse approaches tailored to specific genetic backgrounds. In human pharmacokinetics, the interplay between age and genotype introduces complexities, with advancing age resulting in almost a 1.5-fold increase in systemic exposure to certain drugs, influenced by specific genotypes [70]. Biological sex adds an additional complexity on top of age and genotype. Longevity interventions exhibit robust sex dimorphism, with many interventions demonstrating more pronounced efficacy in males. This observation calls for future studies in geroprotection to actively seek out drugs with positive effects in females, especially in the context of diseases such as AD that have a significant female bias [71]. Our study contributes to the growing understanding that genotype plays a pivotal role in determining the optimized responsiveness of interventions for age-related conditions. This aligns with the overarching concept of personalized medicine, emphasizing the need for tailored approaches that consider individual genetic profiles for therapeutic outcomes.

In summary, we found that 17αE2 improves a range of systemic and neural outcomes in an *APOE*-dependent manner. Specifically, healthspan benefits of 17αE2 were observed more strongly in *APOE4* mice, a genotype that generally associates with poorer mortality and aging outcomes in both mice and humans. Across all measures, *APOE4* mice trended towards worse outcomes relative to *APOE3* mice and generally exhibited stronger benefits from 17αE2 treatment. The ability of 17αE2 to alleviate adverse phenotypes across multiple systems linked to *APOE4* genotype implies that 17αE2 might offer mitigation of the *APOE4*-associated risks including cognitive decline and AD.

### Limitations of the study

There are a few limitations to this study. First, only male mice were studied and thus, the observed *APOE* genotype-dependent effects of 17αE2 may differ in females. As the effects of *APOE4* interact with sex in both human [72, 73] and rodent [74, 75] studies, future research should specifically explore the effects of 17αE2 in female mice with human *APOE3* and *APOE4* despite the absence of significant longevity promotion by 17αE2 in female mice [13]. Second, mouse strain can affect efficacies of longevity promoting interventions [76, 77]. It is noteworthy that the key studies of 17αE2 on longevity employed UM-HET3 mice [13, 29], while mice on a predominantly C57BL/6 background were utilized in this study. Strain-specific variations may reasonably influence the findings observed in *APOE3/4* humanized mice. Lastly, although our study implies a potential protective role of 17αE2 against *APOE4*-associated cognitive decline and AD risk, this research does not directly address AD pathology. Further investigation of 17αE2 effects in the presence of AD-related pathways and factors, such as amyloidosis and tauopathy, should be pursued in additional rodent models to establish a more comprehensive understanding across the complexities of AD in humans.

## Methods

### Materials and methods

#### Animals and treatment

All male mice were homozygous for knock-in of human *APOE3* or *APOE4*. The mice were generated from a breeding colony of EFAD (*APOE*^+/+^, *5xFAD*^+/-^) mice [28], but were non-carriers of the 5xFAD Alzheimer’s-related genes (*APOE*^+/+^, *5xFAD*^-/-^). The colony was started from breeding pairs generously provided by Mary Jo LaDu (University of Illinois at Chicago). Mice were maintained in a vivarium under controlled temperature, a 12:12 light/dark schedule (lights on at 6:00am), group housing when applicable, and *ad libitum* access to water and food (except when specified otherwise). At 10-10.5 months of age, *APOE3* and *APOE4* mice were randomized to two dietary treatment groups (n=17-22/group): TestDiet 5LG6 chow [67.3% carbohydate, 20.5% protein, 12.1% fat] formulated with 0 (Control diet) or 14.4 ppm 17αE2 (Steraloids, Newport, RI) by TestDiet (Richmond, IN). This 17αE2 dosage and delivery method were demonstrated to increase mean and maximum lifespan in male mice [78]. The experimental design is summarized in Figure 1A.

The mice were kept on the diets over a treatment period of 20 weeks, during which body weight and food consumption were recorded weekly. Body composition was measured at weeks 0 and 20 of treatment using a Bruker minispec whole body composition analyzer (Bruker LF90 Minispec, Bruker Optics, Billerica, MA). Mice were monitored daily for overall health, appearance, and euthanasia criteria including >20% body weight decrease, lethargy, and poor grooming. After the 20-week experimental period, mice were euthanized through inhalation of carbon dioxide after overnight food withdrawal, followed by transcardial perfusion with 20mL ice-cold 0.1M PBS. The brains were rapidly removed, and one hemibrain was immersion-fixed for 48 hours in 4% paraformaldehyde/ 0.1M PBS, then stored at 4°C in 0.1M PBS/ 0.03% NaN_3_ until processing for immunohistochemistry. The other hemibrain was dissected into cortex, hippocampus, and hypothalamus. All portions were snap frozen for RNA or protein extraction. Plasma was collected and stored at -80°C. Visceral and retroperitoneal fat pads and livers were dissected and weighed, all were snap frozen. Mice used for RNA-seq and lipidomics were not subjected to behavior, GTT, or fasting to avoid long-lasting effects of stress. After the 20-week experimental period, mice were euthanized through inhalation of carbon dioxide and transcardially perfused with 20mL ice-cold 0.1M PBS. Whole blood was collected via cardiac puncture prior to perfusion for plasma isolation. One hemibrain was immediately used for microglia isolation. The other hemibrain was dissected into cortex, hippocampus, and hypothalamus. All procedures were conducted in accordance with National Institutes of Health guidelines, under the supervision of veterinary staff, and following a protocol (#21269) approved by the University of Southern California Institutional Animal Care and Use Committee.

#### Glucose measurements

Mice were fasted for 16 hours overnight, and blood glucose levels were measured at week 0. At week 19, all mice (except for the RNA-seq cohort) were assessed using a glucose tolerance test (GTT) following 16 hours of fasting. In brief, baseline blood glucose was determined, then animals were orally gavaged with 20% D-glucose in water (2g/kg). Blood glucose levels were measured through tail vein bleed and recorded 15, 30, 60 and 120 minutes after administration of the glucose using the Precision Xtra Blood Glucose and Ketone Monitoring System (9881465; Abbott Laboratories, Abbott Park, IL).

### Behavioral assessments

#### Open Field

A standard open field test was performed at week 15 on all mice (except for the RNA-seq cohort). The mice were moved to the behavior room 30 minutes prior to testing to acclimate. Mice were then placed into an open field box (40cm x 40cm) and allowed to explore freely for 10 mins. The following behaviors were recorded: velocity (cm/s); travel distance (cm); ambulatory time (s).

#### Spontaneous alternation test

At 15 weeks animals were tested for spontaneous alternation behavior in the Y maze, a hippocampus-dependent task that assesses attention to novelty and spatial memory [79, 80]. Mice were brought to the behavior room to acclimate 30 minutes prior to testing. The Y maze consists of 3 arms 40cm long and 7cm wide. Each mouse was placed in one arm and allowed to explore the arena for 5 minutes. Arm entries were recorded if the mouse placed 2 paws into the arm. Three consecutive, nonrepeating entries were considered a correct alternation.

#### Barnes maze

The Barnes maze test was performed at week 16 using a modified Barnes maze protocol [81]. The maze consisted of an open circular platform (91.5cm in diameter) with 20 evenly spaced holes (5cm in diameter) located along the border with a rectangular escape box (11cm L x 5cm W x 5xm H) located beneath one hole. Using spatial-visual clues on each side of the platform, mice were trained to find the escape box. On each day of testing, mice were taken into the behavior room 30 minutes prior to testing to allow them to acclimate. On the first day of the Barnes maze the escape box was removed, and mice were habituated to the maze. Each mouse was given a single habituation trial in which they were placed in an opaque cylinder in the center of the maze. After 10 seconds had elapsed the cylinder was removed, allowing the mice to explore the maze freely for 3 minutes under red light. On the second day of the Barnes maze, mice were placed in a cylinder in the center of the maze, a bright light and buzzer were turned on, and after 10 seconds had elapsed the cylinder was removed. Mice were gently guided into the escape box, after which the light and buzzer were turned off. Mice stayed in the escape box for 1 minute before they were returned to their home cages. After this initial training session, each testing day (including the initial training session day) consisted of 3 training trials per day with an intertrial interval of 15 minutes. Between trials, mice were kept in their own home cages and the maze was cleaned with 70% ethanol. Acquisition training continued for 3 more days (4 training days total). During each trial, mice were placed in a cylinder in the center of maze, a light and buzzer were activated, and after 10 seconds the cylinder was removed. Mice were given 3 minutes to explore freely and locate the escape box. The trial ended either when 3 minutes elapsed, or mice found and entered the escape box. Once a mouse found and entered the escape box the light and buzzer were turned off, and the mouse stayed inside the box for 1 minute. If the mouse did not find the escape box after 3 minutes of exploration, they were gently guided into the escape box and stayed inside the box for 1 minute. To ensure that the hidden escape box was not visible to the mice, three decoy boxes (5cm L x 5cm W x 2.5cm H) were placed throughout the maze. 48 hours after the last acquisition trial, mice were given a probe trial in which the escape box was removed. On the probe trial day, mice were placed in the cylinder in the center of maze and a light and buzzer were turned on. After 10 seconds the cylinder was removed, and mice were given 3 minutes to explore freely. All tests were recorded using Noldus Ethovision XT software version 14.

#### Novel object placement and recognition

Novel object placement (NOP) and recognition (NOR) were performed at week 17 of treatment using a modification of a previously described protocol [82]. After the Barnes maze probe trial, one small plastic Lego block was placed in each home cage to habituate the mice to the objects. On each day of testing, mice were taken into the behavior room to habituate for 30 minutes prior to testing. Forty-eight hours after the introduction of the block, mice were placed in the empty arena (40cm x 40cm), facing the wall that was nearest to the experimenter, and explored for 5 minutes to habituate to the arena. Twenty-four hours after the first habituation, the second habituation was performed in the same way. Twenty-four hours after the second habituation session, mice were placed in the empty arena and explored for 2 minutes, followed by returning to their home cages. The sampling trial consisted of two identical objects placed in the northeast and northwest corners of the arena, and mice were placed in the arena with their heads positioned opposite to the objects. They explored both objects until they accumulated a total of 30 seconds of exploration time, with a maximum of 20 minutes allowed for completion of training. Four hours after sampling, mice were placed in the testing arena in which one of the identical objects was moved to the southeast or southwest corner. The location of the moved object was counter-balanced across all mice. Mice were allowed to explore both objects until they accumulated a total of 30 seconds of exploration time, with a maximum of 20 minutes allowed for completion. Twenty-four hours after sampling, mice were placed in the testing arena in which a novel object was substituted for one of the familiar objects. They explored both objects until they accumulated a total of 30 seconds of exploration time, with a maximum of 20 minutes allowed for completion. Data are presented as discrimination index, which is defined as the time spent with the novel object minus time spent with the familiar object divided by total exploratory time. All tests were recorded using Noldus Ethovision XT software version 14.

#### Frailty measurement

The frailty index (FI) score was calculated for each mouse using a 25-point frailty index, which was modified from a previously described 31-item frailty index [30] to omit measures of auditory deficits. FI assessment included evaluation of the integument, the physical/musculoskeletal system, the ocular/nasal system, the digestive/urogenital system, the respiratory system, and signs of discomfort. The severity of each deficit was rated with a simple scale: 0, 0.5, and 1. A score of 0 was given if there were no signs of a deficit, a score of 0.5 was given to a mild deficit, and a score of 1 means a severe deficit. All these values were summed, giving a frailty score between 0 and 25 for each mouse. The researcher performing the measurement was blinded to all groups.

#### Plasma leptin quantification

Whole blood was collected via cardiac puncture and placed into a plasma collection tube containing EDTA (367856; BD Biosciences, Franklin Lakes, NJ). Plasma leptin was determined using an ELISA kit (EZML-82K; Millipore Sigma, St. Louis, MO) according to the manufacturer’s instructions.

#### Liver Oil Red O (ORO) staining and quantification

Frozen livers were sectioned at 10 µm using a cryostat at -13°C. 4 pieces were collected on the same glass slide for each animal (N=8/group). The slides were stored in the -20°C freezer until ORO staining. Frozen liver sections were brought to room temperature, dipped 10 times into freshly prepared 60% triethyl phosphate, then stained with a solution of 0.5% Oil Red O/60% triethyl phosphate for 16 minutes. Sections were rinsed in a gentle stream of running water for 2 minutes and put into clean water before mounting with prewarmed glycerin jelly. Stained sections were stored at room temperature for one day after which high magnification brightfield images (four sections/liver, one field/section, 20X objective) were collected by unbiased sampling for a total of 4 images per liver. Images were captured using an Olympus BX50 microscope and DP74 camera paired with a computer running CellSens software v1.11(Olympus). Images were converted to grayscale and thresholded using NIH ImageJ 1.50i to yield binary images separating positive and negative immunostaining. The Analyze-Measure tool was used to obtain the value of the percentage of the ORO-positive area.

#### DNA methylation sequencing and epigenetic age calculation

Frozen liver tissue (n=10/group) were sent to Zymo Research for processing through their DNAge Service (Tustin, CA). Briefly, DNA was purified from the frozen liver using the Quick-DNA Miniprep Plus kit (D4068, Zymo Research, Tustin, CA). After quality and quantity checks, bisulfite conversion was performed using the EZ DNA Methylation-Lightening kit (D5030, Zymo Research, Tustin, CA). Samples were enriched for >500 age-associated gene loci and sequenced on an Illumina NovaSeq6000 instrument. Sequenced reads identified by Illumina’s base calling software were aligned to the mouse reference genome using Bismark. Cytosine methylation level was determined as the number of reads reporting a C, divided by the total number of reads reporting a C or T. DNA methylation values were used to assess DNAge according to Zymo’s proprietary DNAge predictor.

#### Lipidomics

Lipidomics was performed by the UCLA Lipidomics Core. For homogenized tissue, 50-100 mg of tissue were collected in a 2mL homogenizer tube pre-loaded with 2.8mm ceramic beads (19-628; Omni, Kennesaw, GA). 0.75mL PBS was added to the tube and homogenized in the Omni Bead Ruptor Elite (3 cycles of 10 seconds at 5 m/s with a 10 second dwell time). Homogenate containing 2-6mg of original tissue was transferred to a glass tube for extraction. A modified Bligh and Dyer extraction [83] was carried out on all samples. Prior to biphasic extraction, an internal standard mixture consisting of 70 lipid standards across 17 subclasses was added to each sample (AB Sciex 5040156, Avanti 330827, Avanti 330830, Avanti 330828, Avanti 791642). Following two successive extractions, pooled organic layers were dried down in a Thermo SpeedVac SPD300DDA using ramp setting 4 at 35°C for 45 minutes with a total run time of 90 minutes. Lipid samples were resuspended in 1:1 methanol/dichloromethane with 10mM Ammonium Acetate and transferred to robovials (Thermo 10800107) for analysis.

Samples were analyzed on the Sciex 5500 with DMS device (Lipidyzer Platform) with an expanded targeted acquisition list consisting of 1450 lipid species across 17 subclasses at the UCLA Lipidomics Core. Differential Mobility Device on Lipidyzer was tuned with EquiSPLASH LIPIDOMIX (Avanti 330731). Data analysis performed on an in-house data analysis platform comparable to the Lipidyzer Workflow Manager [84]. Instrument method including settings, tuning protocol, and MRM list available in [84]. Quantitative values were normalized to mg of tissue.

#### Lipidomics bioinformatics analysis

Lipids not detected across all samples were discarded. A total of 841 lipids were originally detected across plasma and cortex; after filtering, 565 remained for plasma and 501 for cortex. The dataset was first normalized to the amount of tissue or plasma. Then, variance stabilizing normalization was applied to the data using ‘limma’ v.3.48.3, as recommended by previous studies [85, 86] . Differential analysis was performed using ‘limma’ in R, and lipids with an FDR < 5% were considered statistically significant. Lipid ontology enrichment analysis was performed using the LION web-based ontology enrichment tool with all detected lipids used as the background [32].

#### Microglia isolation from fresh mouse brain

Following dissection, one hemibrain from a dedicated cohort that included all experimental groups (n= 4-5/group) was used to isolate microglia for RNA sequencing. This hemibrain was temporarily stored in 5mL of HBSS buffer (w/o Ca^2+^, Mg^2+^) (88284, Thermo Fisher Scientific, Waltham, MA). Tissue dissociation was performed using the Worthington Papain Dissociation System (LK003150; Worthington Biochemical, Lakewood, NJ) according to the manufacturer’s instructions. The dissociated cell pellet was resuspended in 1mL of MACS buffer (130-091-221; Miltenyi Biotec, Bergisch Gladback, North Rhine-Westphalia, Germany). Microglia were isolated using Miltenyi CD11b Microglia Microbeads (130-093-636; Miltenyi Biotec, Bergisch Gladback, North Rhine-Westphalia, Germany) according to the manufacturer’s instructions. Cell number and viability were determined using trypan blue exclusion on an automated COUNTESS cell counter (Thermo Fisher Scientific, Waltham, MA). Purified cells were pelleted at 300x*g* then snap-frozen in liquid nitrogen until RNA extraction.

#### RNA isolation and RNA-seq library preparation

For RNA isolation, frozen cell pellets were resuspended in 1 ml of Trizol reagent (15596018; Invitrogen, Carlsbad, CA) and total RNA was purified following the manufacturer’s instructions. RNA quality was assessed using the Agilent Bioanalyzer platform at the USC Genome Core using the RNA integrity number. Then, 500 ng of total RNA was subjected to ribosomal-RNA depletion using the NEBNext rRNA Depletion kit (E7850L; New England Biolabs, Ipswich, MA), according to the manufacturer’s protocol. Strand specific RNA-seq libraries were then constructed using the SMARTer Stranded RNA-seq kit (634485; Clontech, Kusatsu, Shiga, Japan), according to the manufacturer’s protocol. Libraries were quality controlled on the Agilent Bioanalyzer 2100 platform at the USC Genome Core before multiplexing the libraries for sequencing. Paired-end 150-bp reads were generated on the Illumina NovaSeq6000 platform at the Novogene Corporation (Sacramento, CA). Raw sequencing reads have been deposited to the Sequence Read Archive under accession PRJNA1078754.

#### RNA-seq bioinformatic analysis pipeline

Paired-end 150-bp reads were hard-trimmed to 75 bp using Trimmomatic v0.39 [87]. Trimmed reads were mapped to the mm39 genome reference using STAR v.2.7.0e [88]. Read counts were assigned to genes from the UCSC mm39 reference using subread v.2.0.2 [89] and were imported into R version 1.4.1717 to perform differential gene expression analysis.

Only genes with mapped reads in at least half of the RNA-seq libraries were considered to be expressed and retained for downstream analysis. We used surrogate variable analysis (sva) to estimate and correct for unwanted experimental noise [90]. R package ‘sva’ v.3.40 was used to estimate surrogate variables and the removeBatchEffect function from ‘limma’ v.3.48.3 was used to regress out the effects of surrogate variables and RNA integrity differences (RNA integrity number scores) from raw read counts. The ‘DESeq2’ R package (DESeq2 v.1.32.0) was used for further processing of the RNA-seq data in R [91]. Genes with FDR < 5% were considered statistically significant and are reported in Supplementary Table 2. We found a non-linear relationship of treatment between the genotypes and thus modeled treatment and genotype separately. The following comparisons were performed: *APOE3* control to *APOE4* control, *APOE3* control to *APOE3* 17αE2, *APOE4* control to *APOE4* 17αE2, and *APOE3* control to *APOE4* 17αE2.

#### Dimensionality reduction

To perform multi-dimensional scaling (MDS) analysis [46], we used a distance metric between samples based on the Spearman’s rank correlation value (1-Rho), which was then provided to the core R command ‘cmdscale’. Dimensionality reduction was applied to DESeq2 VST-normalized counts.

#### Functional enrichment analysis

The Gene Set Enrichment Analysis (GSEA) paradigm through its implementation in the R package ‘ClusterProfiler’ v4.0.5 [92], and Bioconductor annotation package ‘org.Mm.eg.db’ v3.13.0 were used to perform the functional enrichment analysis. The DEseq2 t-statistic was used to generate the ranked list of genes for functional enrichment analysis, for both genotype and treatment effects. The top 5 up- and down-regulated Gene Ontology (GO) terms per genotype are shown in figures if at least that many passed the FDR < 5% significance threshold (Figure 5G,H). All significant GO terms are reported in Supplementary Table 2.

#### Immunohistochemistry

Hemibrains fixed in 4% paraformaldehyde (A11313, Alfa Aesar, Ward Hill, MA) for 48 hours were transferred into 20% sucrose in PBS overnight at 4°C until they sank to the bottom of the tube. Brains were then coronally sectioned at 20μM using a cryostat (Leica Biosystems, Deer Park, IL). Four sections per well were stored in 0.03% sodium azide in PBS at 4°C until immunohistochemistry was performed. Every eighth section from approximately -0.95mm to - 2.90mm were immunostained with ionized calcium binding adaptor molecule 1 (Iba-1; 1:2000; 019-19741; FUJIFILM Wako, Richmond, VA) or glial fibrillar acidic protein (GFAP; 1:1000; G3893-100uL; Dako, Santa Clara, CA). For Iba1 staining, brain sections were first incubated with 10nM EDTA (pH = 6) at 95°C for 10 minutes. No antigen retrieval pretreatment was performed for GFAP staining. For both Iba1 and GFAP staining, sections were rinsed with 0.1M Tris-buffered saline (TBS) and treated with an endogenous peroxidase blocking solution (3% H_2_O_2_, 10% methanol in TBS) for 10 minutes. Sections were rinsed in 0.2% Triton-X in TBS before being blocked for 30 minutes in blocking solution. For Iba1, the blocking solution contained 2% bovine serum albumin (BSA) in TBS. For GFAP, the blocking solution contained 2% BSA and 2% normal goat serum in TBS. Sections were incubated with primary antibodies in their respective blocking solutions overnight at 4°C. The next day, sections were rinsed in 0.1% Triton-X in TBS and incubated for 1 hour in their respective biotinylated secondary antibody diluted in blocking solution. Sections were rinsed once more in 0.1% Triton-X in TBS then incubated in an avidin-biotin complex (PK-6100; Vectastain ABC Elite kit, Vector Laboratories, Newark, CA) for 1 hour. To visualize immunoreactivity, sections were incubated for 5 minutes using diaminobenzidine tetrahydrochloride (SK-4100; Vector Laboratories, Newark, CA).

#### Iba1 and GFAP Load

To quantify Iba1 and GFAP immunoreactivity, non-overlapping high magnification brightfield images were collected from the CA1 hippocampal subfield (three fields/section, 20x objective) across four tissue sections per brain, for a total of ∼12 images per brain. Images were captured using an Olympus BX50 microscope and DP74 camera paired with a computer running CellSens software v1.11(Olympus). Images were converted to grayscale and thresholded using NIH ImageJ 1.50i to yield binary images separating positive and negative immunostaining. Iba1 and GFAP load was calculated as the percentage of total pixels that was positively immunolabeled.

#### Protein extraction

Brain cortices were homogenized with a motorized pestle in RIPA buffer without SDS (30 mg tissue: 150 μL) with protease (P2714, Millipore, Bedford, MA, USA) and phosphatase (78427, Thermo Fisher Scientific, Waltham, MA) inhibitors as previously described [61] . Homogenates were centrifuged at 10,000 × g for 1 hour at 4°C, and supernatant was recovered for evaluation of soluble amyloid peptides. Proteins were quantified using Pierce’s 660nm assay (22660, Thermo Fisher Scientific, Waltham, MA).

#### Lipid Rafts

Lipid rafts were isolated by kit using 35mg of cortex (LR-039, Invent Biotechnologies, Plymouth, MN). Lipid rafts were previously validated against traditional ultracentrifugation methods [61].

#### Dot blots

25 μg of RIPA or 5ug of lipid raft protein lysate was loaded onto a dot blot apparatus (Bio-Rad, Hercules, CA, USA) and were filtered through 0.45μm PVDF for 2 hours by gravity filtration. Membranes were stained with Revert 700 (926-11011, LICOR, Lincoln, NE, USA), imaged, and blocked for 1 hour with Intercept blocking buffer (927-70001, LICOR) before incubation for 16 hours with primary antibodies for Aβ40, (Biolegend, San Diego, CA, USA), and HNE (ABN249, Millipore, Bedford, MA, USA). Membranes were visualized on a LICOR 9120 using fluorescent-conjugated secondary antibodies. Images were analyzed by ImageJ and corrected by total protein load.

#### Statistics

All data are reported as the mean ± the standard error of the mean. Data were analyzed using GraphPad Prism version 5 (biochemical, behavioral, and metabolic data) or R version 1.4.1717 (‘omic’ data). All data were checked for normal distribution using the Shapiro-Wilk test. If a dataset was found not normally distributed, Mann-Whitney tests were used. Non-normally distributed datasets include oil red o staining, spontaneous alternation behavior, novel object placement, Iba1 immunohistochemistry, and open field: time in center. For normally distributed data, which described most of the group comparisons, two-way or three-way ANOVAs followed by Tukey post-hoc were performed. Two-way repeated measure analysis of variance followed by Tukey post tests were run for all data measured over time. Comparisons with *p* < 0.05 were considered statistically significant.

## Data and code availability

All sequencing data were deposited to SRA under accession PRJNA1078754. Lipidomics (DOI:10.6084/m9.figshare.25346143) and DNAge (DOI:10.6084/m9.figshare.25346143) data were deposited to FigShare. All R scripts are available on Github repository https://github.com/BenayounLaboratory/17aE2_APOE.

## Author Contributions

C.J.P., B.A.B., and C.E.F. designed the study. C.J.M, A.C., W.Q., M.T., J.G.L., S.N., and B.B. performed experiments. C.J.M performed data analyses. C.J.M performed bioinformatics analysis, and O.S.W. independently checked the code. C.J.M., B.A.B. and C.J.P. wrote the manuscript. All authors approved the final version of the manuscript.

## Acknowledgements

This study was supported by a grant from the Cure Alzheimer’s Fund (C.J.P., C.E.F., B.A.B.). C.J.M. was supported by NIH/NIA grants T32 AG052374 (S.P. Curran) and F31 AG084279 (C.J.M.). The authors thank the UCLA Lipidomics Laboratory for their contributions and Ms. Tavia Roache, Ms. Bayla Breningstall and Ms. Tyne McHugh for their experimental contributions. Some images from schematics obtained from Freepik.

