## Supplementary Figure 1 for "Protection against *APOE4*-associated aging phenotypes with the longevity-promoting intervention 17α-estradiol in male mice"

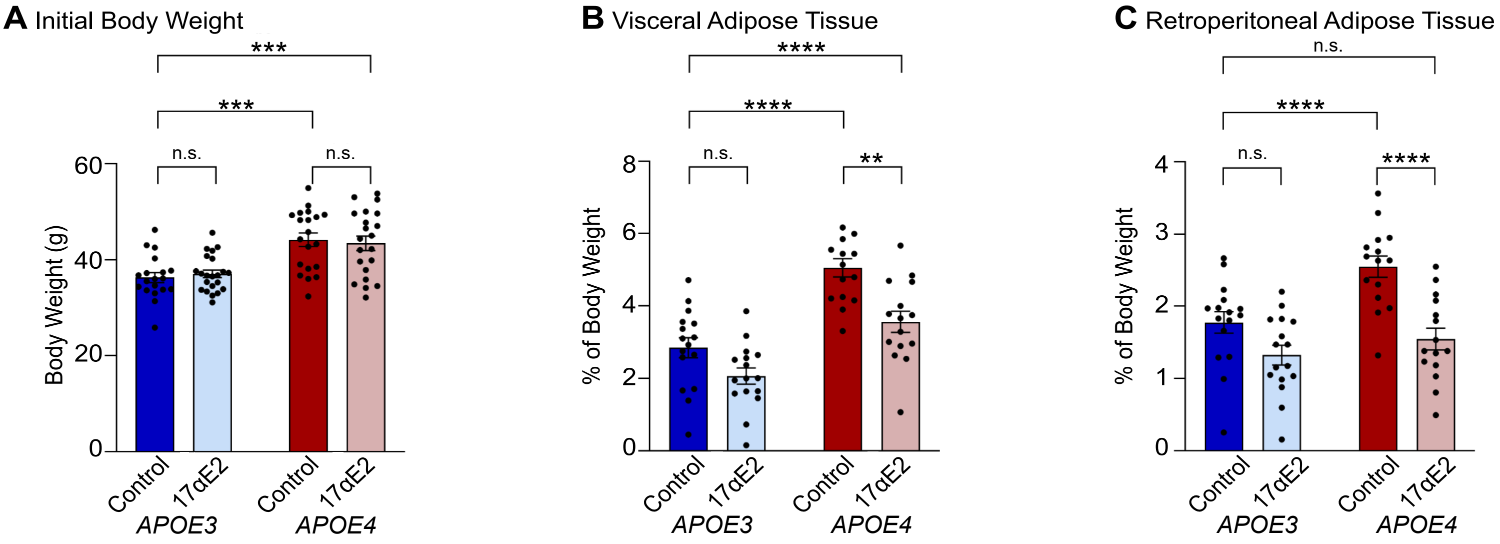


**Supplemental Figure 1. APOE4-related adiposity is decreased with 17αE2 treatment. (A) Starting body weights of APOE3 and APOE4 male mice prior to treatment initiation (n=19-22/group). (B) % of visceral adipose tissue compared to overall body weight after 20 weeks of 0 or 14.4ppm 17αE2 (n=15-16/group). (C) % of retroperitoneal adipose tissue compared to overall body weight after 20 weeks of 0 or 14.4ppm 17αE2 (n=15-16/group). In (A) through (C), dark blue indicates APOE3 control, light blue indicates APOE3 17αE2, dark red indicates APOE4 control, and light red indicates APOE4 17αE2. Data show mean ± SEM. Asterisks denote statistical significance: ** $p < 0.01$, *** $p < 0.001$, **** $p < 0.0001$ in 2-way ANOVA Tukey post-hoc test.**
