## Supplementary Figure 2 for "Protection against *APOE4*-associated aging phenotypes with the longevity-promoting intervention 17α-estradiol in male mice"

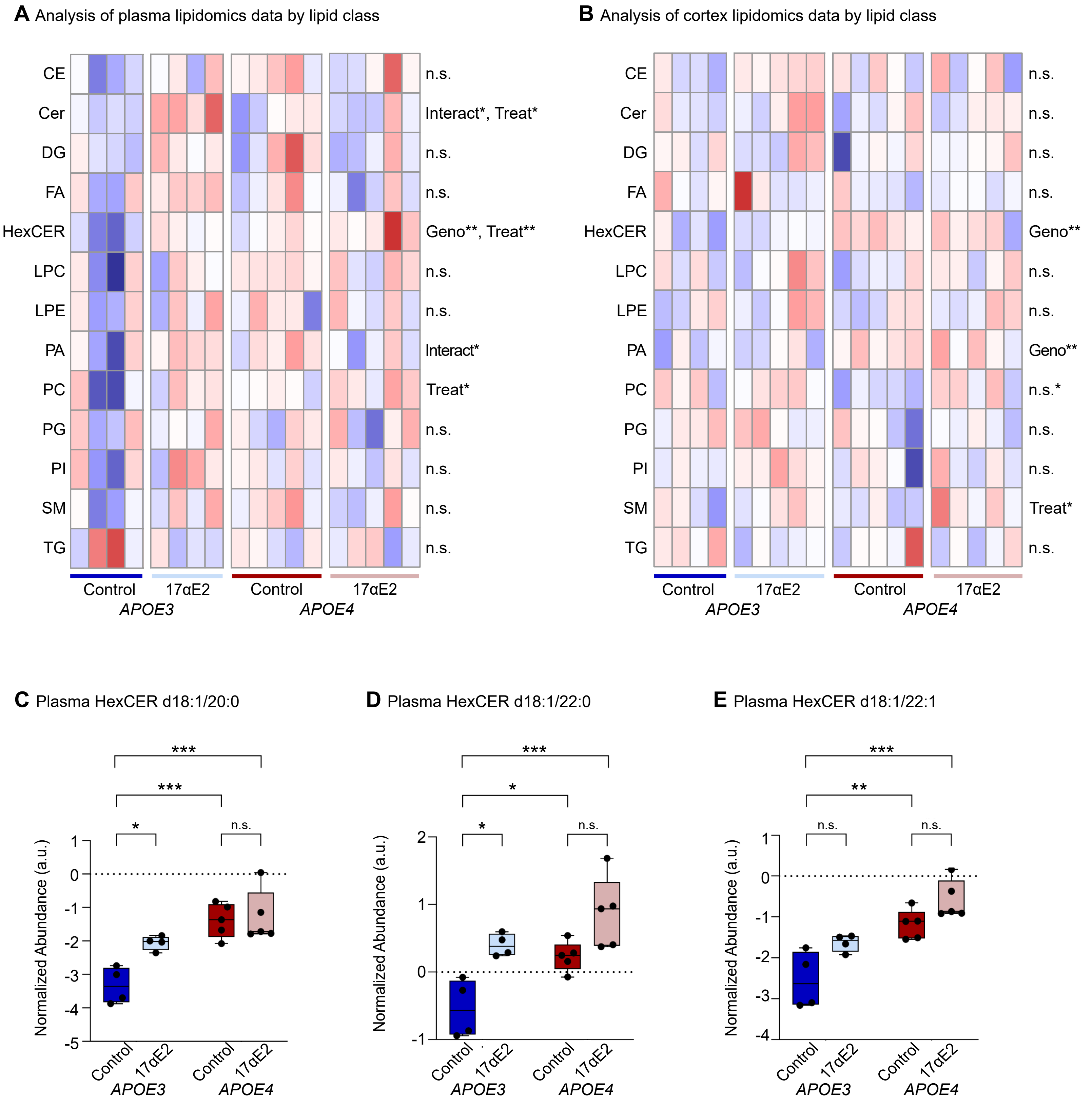


**Supplemental Figure 2. Plasma lipidomics show genotype and treatment differences. (A) Analysis of plasma lipidomics by lipid class (n=4-5/group). Data shows Cholesterol esters (CE), Ceramides (Cer), Diacylglycerols (DG), Free Fatty Acids (FA), Hexosyl ceramides (HexCER), lysophosphatidylcholine (LPC), lysophosphatidylethanolamine (LPE), Phosphatidic acid (PA), Phosphatidylcholine (PC), Phosphatidylglycerol (PG), Phosphatidylinositol (PI), Sphingomyelin (SM), and Triacylglycerols (TG). (B) Analysis of cortex lipidomics by lipid class. Data shows identical lipids found in (A) (n=4-5/group). (C) Plasma HexCER d18:1/20:0 levels (n=4-5/group). (D) Plasma HexCER d18:1/22:0 levels (n=4-5/group). (E) Plasma HexCER d18:1/22:1 levels (n=4-5/group). In (C) through (E), dark blue indicates APOE3 control, light blue indicates APOE3 17αE2, dark red indicates APOE4 control, and light red indicates APOE4 17αE2. Data show mean ± SEM. Asterisks denote statistical significance: * $p < 0.05$, ** $p < 0.01$, *** $p < 0.001$ in 2-way ANOVA Tukey post-hoc test.**
