## Supplementary Figure 3 for "Protection against *APOE4*-associated aging phenotypes with the longevity-promoting intervention 17α-estradiol in male mice"

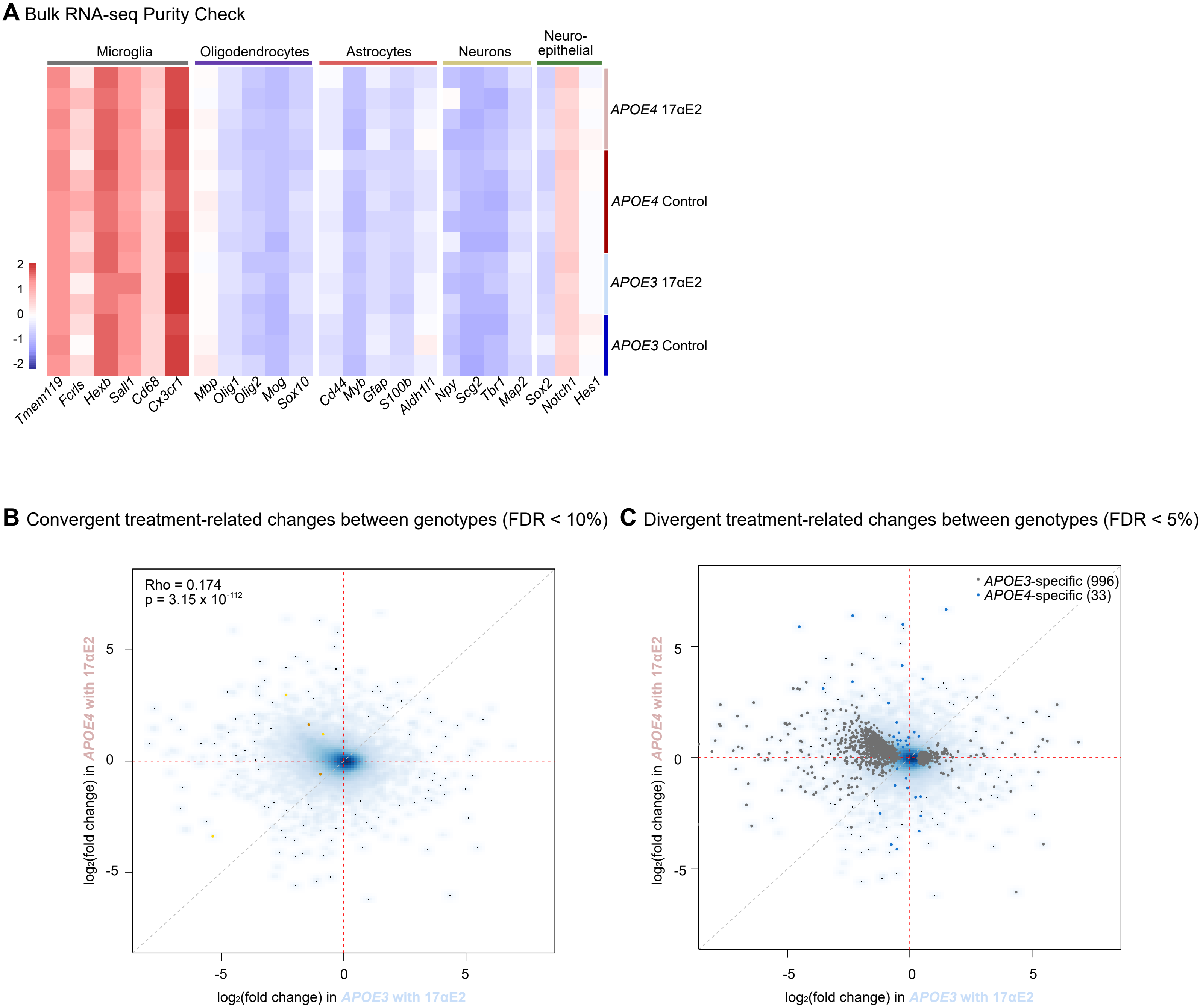


**Supplemental Figure 3. Microglia Bulk RNA-seq. (A) Purity check using gene markers of various brain cell types (n=3-5/group). (B-C) Correlation plot of treatment-related gene expression changes according to DESeq2 in APOE3 vs APOE4 microglia from RNA-seq, showing genes with (B) significant and concordant treatment-regulation in both genotypes (FDR 10% in gold, FDR 5% in blue), or (C) genes with divergent treatment-regulation between genotypes (FDR 5%).**
