## Supplementary Figure 4 for "Protection against *APOE4*-associated aging phenotypes with the longevity-promoting intervention 17α-estradiol in male mice"

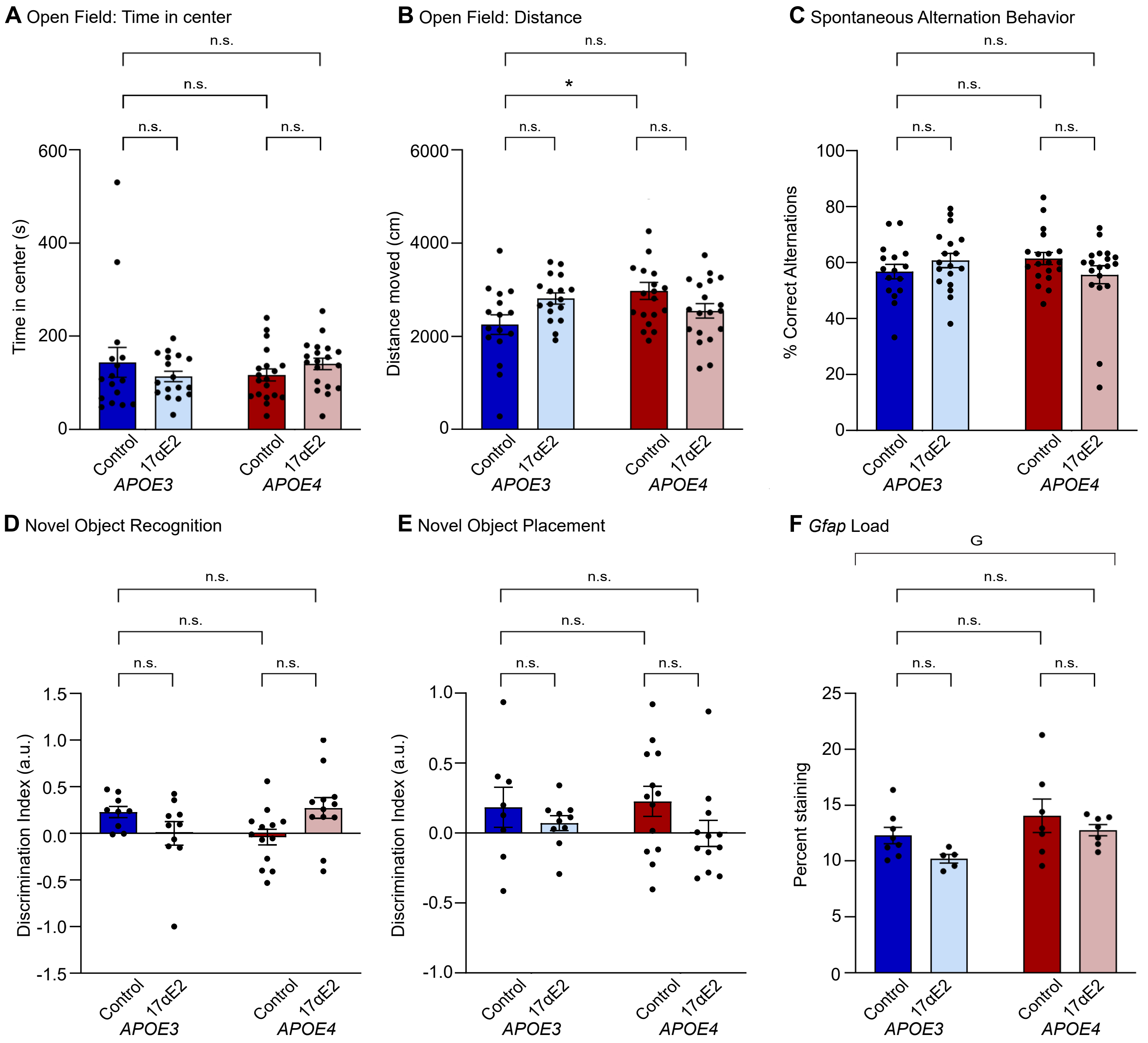


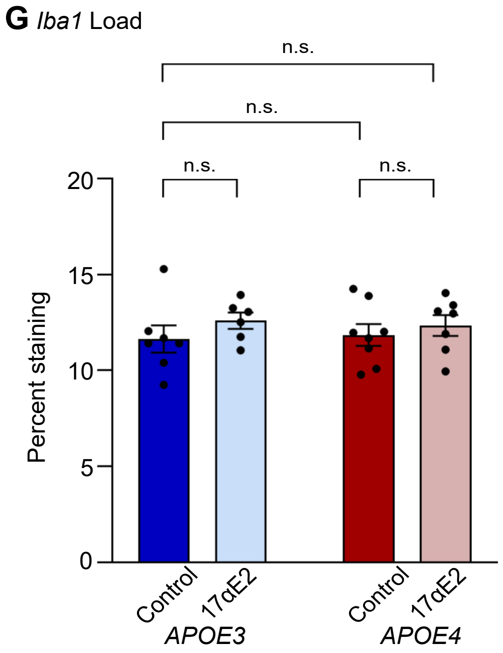


**Supplemental Figure 4. APOE4 mice experience greater neural benefits from 17αE2 compared to APOE3. (A) Time in center of the open field test (n=16-20/group). (B) Total distance traveled in the 10 minutes of the open field test (n=16-20/group). (C) Percent of correct alternations in the spontaneous alternation behavior test (n=16-20/group). (D) Discrimination index of the novel object recognition test (n=9-13/group). (E) Discrimination index of the novel object placement test (n=8-13/group). (F) Immunohistochemical load of the astrocyte marker GFAP in the CA1 region of the hippocampus (n=5-7/group). (G) Immunohistochemical load of the microglial marker Iba-1 in the CA1 region of the hippocampus (n=6-8/group). In (A) through (G), dark blue indicates APOE3 control, light blue indicates APOE3 17αE2, dark red indicates APOE4 control, and light red indicates APOE4 17αE2. Data show mean ± SEM. Asterisks denote statistical significance: * $p < 0.05$ in 2-way ANOVA Tukey post-hoc test.**
