## Supplementary Tables for "Protection against *APOE4*-associated aging phenotypes with the longevity-promoting intervention 17α-estradiol in male mice"

| **Supplementary Table S1. Plasma lipidomics and enrichment analyses.** | | |
| --- | --- | --- |
|  | **A** | Limma significantly regulated lipids between control *APOE3* and control *APOE4* plasma by lipidomics (FDR <5%) |
|  | **B** | Limma significantly regulated lipids between control *APOE3* and 17α-estradiol treated *APOE4* plasma by lipidomics (FDR <5%) |
|  | **C** | LION functional enrichment analysis for differential lipids between control *APOE3* and *APOE4* plasma higher in *APOE3* |
|  | **D** | LION functional enrichment analysis for differential lipids between control *APOE3* and *APOE4* plasma higher in *APOE4* |

**Supplementary Table S2. Microglia bulk RNA-seq and enrichment analyses.**

| **A** | DEseq2 significantly regulated genes between control *APOE3* and control *APOE4* microglia by RNA-seq (FDR <5%) |
| --- | --- |
| **B** | DEseq2 significantly regulated genes between control *APOE3* and 17 alpha-estradiol treated *APOE4* microglia by RNA-seq (FDR <5%) |
| **C** | DEseq2 significantly regulated genes between control *APOE3* and 17 alpha-estradiol treated *APOE3* microglia by RNA-seq (FDR <5%) |
| **D** | DEseq2 significantly regulated genes between control *APOE4* and 17 alpha-estradiol treated microglia by RNA-seq (FDR <5%) |
| **E** | GSEA Gene Ontology (FDR <5%) for differential regulation between control *APOE3* and control *APOE4* microglia by RNA-seq |
| **F** | GSEA Gene Ontology (FDR <5%) for differential regulation between control *APOE3* and 17 alpha-estradiol treated *APOE4* microglia by RNA-seq |
| **G** | GSEA Gene Ontology (FDR <5%) for differential regulation between control *APOE3* and 17 alpha-estradiol treated *APOE3* microglia by RNA-seq |
| **H** | GSEA Gene Ontology (FDR <5%) for differential regulation between control *APOE4* and 17 alpha-estradiol treated *APOE3* microglia by RNA-seq |
